# Atomistic Simulation of Voltage Activation of a Truncated BK Channel

**DOI:** 10.1101/2025.01.08.631907

**Authors:** Zhiguang Jia, Jianhan Chen

## Abstract

Voltage-dependence gating of ion channels underlies numerous physiological and pathophysiological processes, and disruption of normal voltage gating is the cause of many channelopathies. Here, long timescale atomistic simulations were performed to directly probe voltage-induced gating transitions of the big potassium (BK) channels, where the voltage sensor domain (VSD) movement has been suggested to be distinct from that of canonical Kv channels but remains poorly understood. Using a Core-MT construct without the gating ring, multiple voltage activation transitions were observed at 750 mV, allowing detailed analysis of the activated state of BK VSD and key mechanistic features. Even though the S4 helix remains the principal voltage sensor in BK, its vertical displacement is only ∼3 Å and accompanied by significant lateral movements. The nature of the predicted VSD movement is in strong agreement with recent Cryo-EM structural studies of mutant BK channels with constitutively activated VSD. Free energy analysis based on the predicted activation transition yielded a total gating charge of 0.44 *e* per VSD, consistent with the experimental range of 0.48 – 0.65 *e*. We further show that the ability of modest physical movements with a small total gating charge to drive effective voltage gating of BK can be attributed to large gradients in the local electric field as reshaped by the protein. Furthermore, the S4 movement is coupled to the pore opening through a non-canonical pathway that involves the tightly packed S4-S5-S6 interface. These distinct mechanistic features may be relevant to voltage gating of other ion channels where VSDs are not domain-swapped with respect to the pore-gate domain.

**Significance Statement:** The big potassium (BK) channel is the only potassium channel that integrates intracellular calcium signaling with membrane depolarization. It is considered one of the most important channels in cardiovascular and neurological disorders. It has been known that voltage gating of BK channels has distinct features compared to the canonical voltage gating mechanism established through studies of voltage-gated potassium (Kv) channels. Yet, little is known about the molecular nature of voltage sensing and pore activation of BK channels at present. Our work reports the first successful direct simulation of voltage-dependent activation of the big potassium (BK) channel, revealing novel voltage sensing and sensor-pore coupling mechanisms that have largely eluded the community until now.

## Introduction

Voltage-dependent gating of ion channels mediates numerous physiological and pathophysiological processes (*1–5*), by regulating ion flows in response to membrane potential changes in excitable cells such as cardiac myocytes and smooth muscle cells in the lung and blood vessels (*6–11*). Disruption of normal voltage gating is involved in many cardiovascular and neurological diseases, including epilepsy, mental retardation, chronical pain, hypertension, arrhythmias, stroke, and ischemia (*12–20*). Among the family of potassium channels, the large-conductance potassium (BK) channels stand out in several ways (*21–27*). It has the largest single-channel conductance (up to ∼300 ps) and is the only known K^+^ channel to be activated by both intracellular Ca^2+^ and membrane potential. Functional BK channels are homo-tetramers, with each subunit consisting of a seven-helix transmembrane domain (TMD) and a C-terminal Ca^2+^-sensing cytosolic domain (CTD) (Figure 1a). The deactivated state of BK channels contain a physically open central pore (*28–30*), lacking the classical bundle crossing constriction at the intracellular entrance observed in the voltage-dependent potassium (Kv) channels (*31*). Instead, BK channels likely follow the hydrophobic gating mechanism, where the pore undergoes hydrophobic dewetting transition to create a vapor barrier to block ion permeation in the deactivated state (*32–36*). Furthermore, the voltage sensor domains (VSDs) of BK channels are not domain-swapped with respect to the pore-gate domain (PGD), in contrast to the domain-swapped TMD organization of Kv channels (Figure 1b-c). The difference in TMD organization is likely a key factor underlying important but still poorly understood differences in the voltage sensing and gating mechanisms of BK channels (*37–41*) in comparison to the “canonical” framework established mainly through the study of Kv channels (*42–50*).

**Figure 1.**
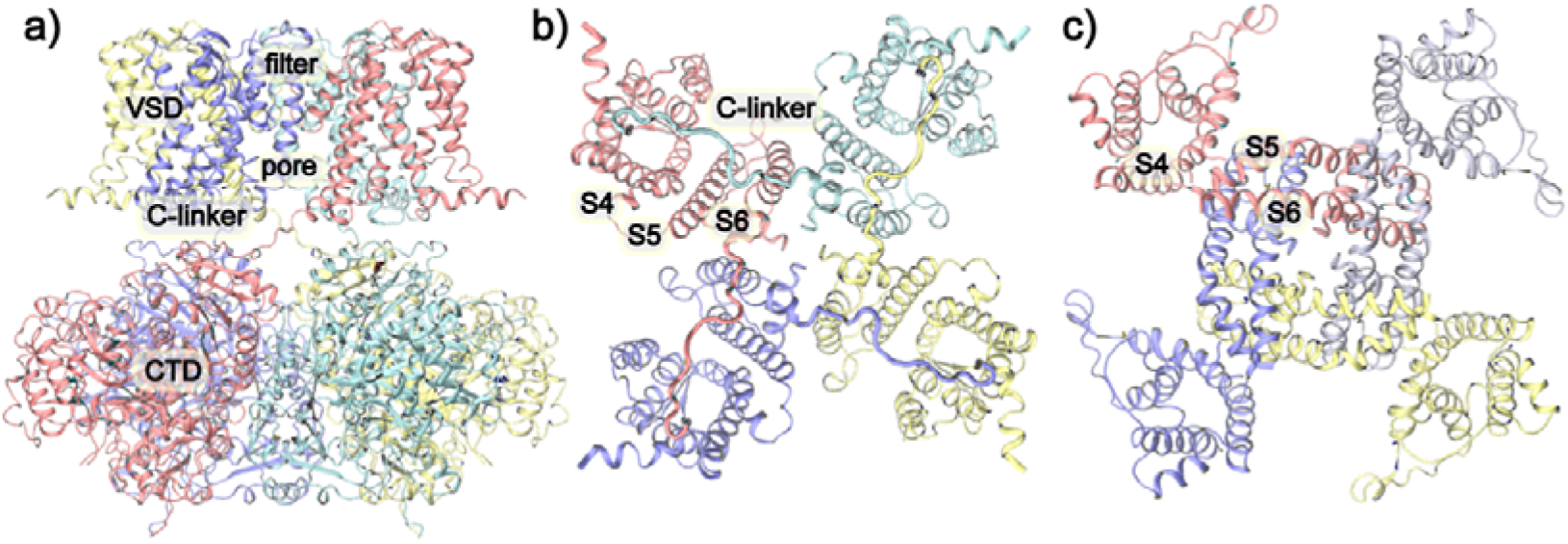
Overall structures of BK and Kv channels. **a**) The overall structure of BK channels in the Ca^2+^-free state (PDB: 6v3g) with key domains and regions labelled. Each monomer is shown in the same color. **b**) Top view of the TMD of BK channels, showing the non-domain-swapped VSD/PGD configuration. **c**) Top view of the TMD of Kv 1.2 channel (PDB: 3lut), where the VSDs and PGDs are domain-swapped. Note the much tighter packing of S4 from VSD and S5/S6 from PGD in BK channels.

In Kv channels, the linker connecting transmembrane (TM) helices S4 and S5 forms a α-helix (∼15 aa) that wraps around the S6, with the S4 of one subunit interacting with the S5 of a neighboring subunit and simultaneously connecting to the S5/S6 within the same subunit through the S4-5 linker. Conversely, in BK channels, the S4-S5 linker is a short loop (∼5 aa), and the S4 solely interacts with S5 within the same subunit (Figure 1). Within the canonical voltage activation of Kv channels, each of the first four Arginine residues in the VSD S4 accounts for about 1*e* gating charge (*51–54*), giving rise to a total gating charge of ∼ 3 - 3.5 *e* per monomer. Membrane depolarization drives S4 to move upward by ∼8 Å, which causes the S4-S5 linker helix to pivot upwards and change the interactions with S6 and releases the S6 helix bundle crossing to physically open the gate (*55–57*). For BK channels, despite sharing many conserved charged residues, the total number of gating charges is only 0.6 *e* per VSD (*22, 58–62*). In addition to Arginine residues on BK S4 (R207, R210 and R213), other residues also appear to contribute to voltage sensing, such as D153 and R167 on S2, D186 on S3 and E219 on S4 (*39, 63, 64*). Furthermore, fluorophore quenching suggested that the movement of BK VSDs during voltage gating was smaller and involved both vertical and lateral motions (*65–67*). At present, there remains significant ambiguity in the identity of gating charges, details of VSD motions, and how they drive the pore opening in BK channels.

In this study, extensive atomistic molecular dynamic (MD) simulations up to 10 μs in length were performed in explicit solvent and membrane to directly probe the voltage-driven activation of BK channels in the Ca^2+^-free state, using the Core-MT construct that does not include the CTD but retains voltage gating (*68, 69*). We were able to directly observe spontaneous opening transitions and ion conductance of the channel under 750 mV membrane voltage within the 10 μs simulation timescale. The observed VSD movements appear to be highly consistent with two recent Cryo-EM structural studies of mutant BK channels with constitutively activated VSDs (*38, 41*). Free energy analysis was preformed to further quantify the total gating charge as well as the contributions of key charged residues. The results are in quantitative agreement with available experimental data (*39, 63, 64*), providing further support for the predicted voltage sensing and gating mechanism. The analysis reveals central roles of voltage-induced displacement of R210 and R213 on S4 in voltage sensing and the strong interactions at the S4-S5-S6 interface in VSD-pore coupling. This mechanism was further validated using steered MD simulations showing that pulling on the charged tip of R210 and R213 alone could ready drive pore opening in Core-MT BK channels. Taken together, the current work provides for the first time a reliable detailed molecular mechanism of BK voltage activation, which is distinct from the canonical voltage gating mechanism of Kv channels. These novel voltage gating principles are likely relevant in understanding the gating and regulation of other non-domain swapping ion channels.

## Results and Discussion

### Direct atomistic simulations of voltage activation of BK channels

Starting from a fully equilibrated closed state, multiple atomistic simulations were performed at 0 and 750 mV membrane voltages for up to 10 μs to directly probe voltage-driven activation of Core-MT BK channels using the special purposed supercomputer Anton 2 (*70, 71*) (see Methods; Figure S1). The simulated voltage of 750 mV is higher than the experimental conditions, where V_1/2_ of Core-MT BK channels is ∼ 244 mV compared to ∼180 mV for the full-length BK channel (*69*), to accelerate protein conformational transitions. Similar voltages have been used in atomistic simulations and found to generate realistic transitions (*56, 72*). We compare the membrane thickness at 300 and 750 mV and the results reveal no significant difference in the membrane thickness (Figure S2). At 0 mV, the Core-MT BK channel remained highly stable in an apparently closed state, with the pore fully dehydrated and not permeable to ions (Figure S1, top row). Similar to what has been observed previously in simulations and cryo-EM maps (*30, 32–35, 38*), lipid tails can enter the dewetted pore through the fenestration gap between the pore lining S6 helices. They are highly dynamic and likely contribute to the stability of the dewetted state of the pore. Note that the absence of CTDs increases the flexibility of pore-lining S6 helices, and the fully relaxed pore profile (red trace in Figure S1d, top row) shows substantial differences compared to that of the Ca^2+^-free Cryo-EM structure of the full-length channel (black trace). Importantly, the VSDs were highly stable and showed minimal movements (∼1 Å or less) for all charged groups as well as the TM helices themselves.

At 750 mV, significant movements were observed with multiple charged groups on S4 (Figure 2a and Figure S1). In particular, the guanidium groups of R210 and R213 moved upward along the membrane normal (z-axis) by up to ∼8 and ∼10 Å, respectively, plateauing after ∼5 μs (e.g., see Figure 2a). Other VSD charges showed much smaller z-displacements. The apparent modest movements of ∼3-4 Å of E219 and R207 seem to be mainly a result of the upshift of S4 itself, which reached a maximum of ∼3 Å during the second half of the simulation (Figure 2b). In contrast, other VSD helices S1-S3 exhibited much smaller z-axis movements (∼1 Å). As illustrated in supplementary Movie S1, the large charge displacements of R210 and R213 charges with modest S4 movement were enabled by side chain snorkeling.

**Figure 2.**
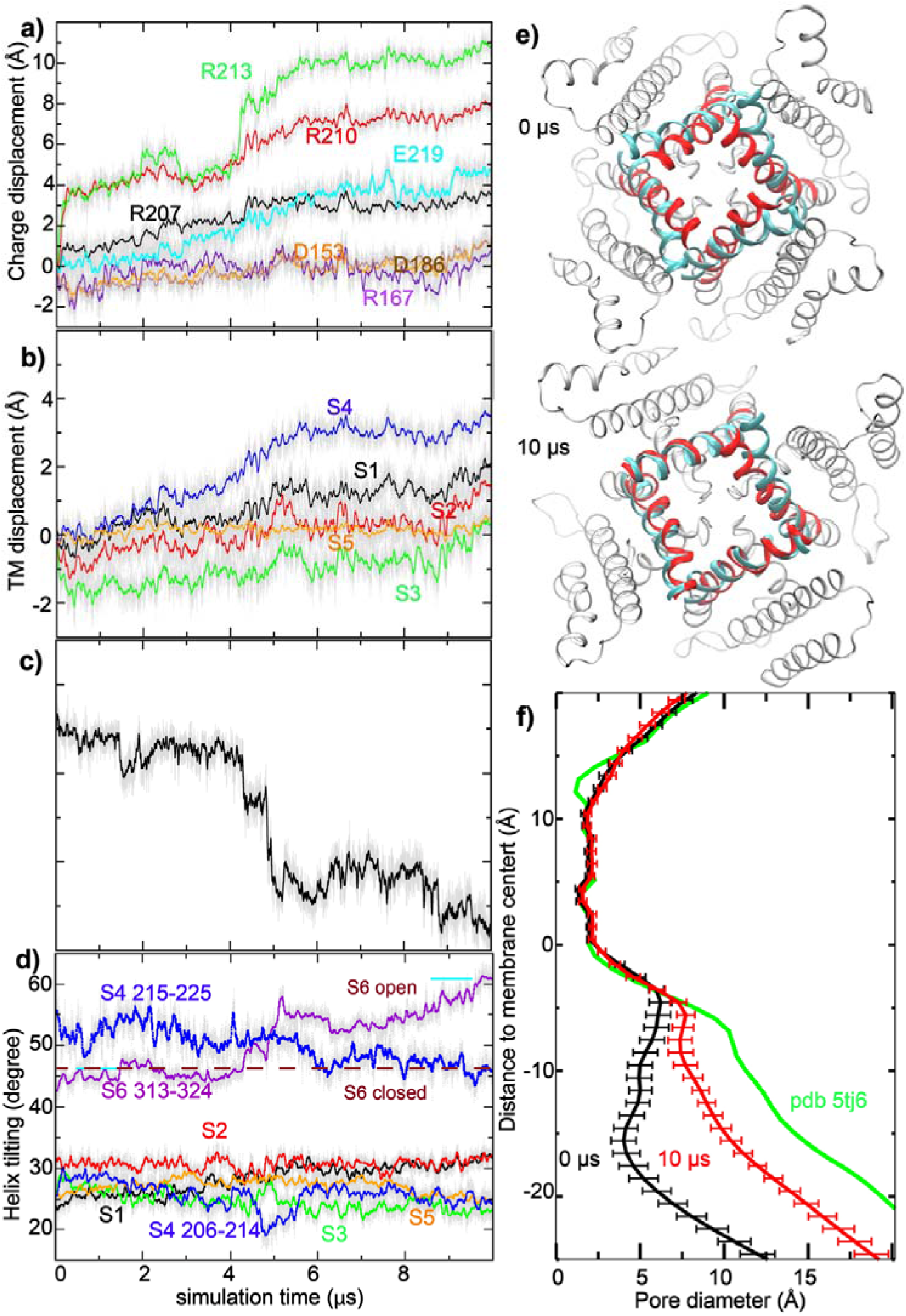
Voltage activation of Core-MT BK channels. **a-d)** Results from a 10-_μ_s simulation under 750 mV (*sim2b* in Table S1). Each data point represents the average of four subunits for a given snapshot (thin grey lines), and the colored thick lines plot the running average. a) z-displacement of key side chain charged groups from initial positions, b) z-displacement of centers-of-mass of VSD helices from initial positions, c) backbone RMSD of the pore-lining S6 (F307-L325) to the open state, and d) tilt angles of all TM helices. The locations of charged groups were taken as those of guanidinium CZ atoms (for Arg) and sidechain carboxyl carbons (for Asp/Glu). Only residues 313-324 of S6 were include in tilt angle calculation, and the values in the open and closed Cryo-EM structures are marked using purple dashed lines for reference in panel d. **e)** Superimposition of the initial (0 _μ_s) and final (10 _μ_s) structures of the pore (residues F307-L325; red cartoon), in comparison to the open Cryo-EM structure (cyan cartoon). The view shown is from the bottom (cytosolic side). **f)** Average pore profiles calculated from the first and last 0.1 _μ_s of *sim2b*, with error bars showing standard error. The pore profile derived from PDB 5tj6 (open state) is shown as a reference.

Voltage-driven VSD movements were apparently coupled with an opening transition of the pore, allowing one to directly observe Core-MT activation and K^+^ conductance during both 10-μs atomistic simulations on Anton 2 (Figure S1). As illustrated in Figure 2 and Movie S2, the backbone root-mean-squared distance (RMSD) of the pore from the open state decreased sharply from ∼4.5 Å around the 4 μs mark to below ∼2.7 Å around the 5 μs mark, as the S4 helix shifted up along z-axis by ∼ 2 Å during the same time span. The pore opening transition mainly involved the increase of the tilt of S6 segments below the glycine hinge (residues 313-324) (Figure 2d). The pore continued to move closer to the open state after the initial rapid response to VSD activation, eventually reaching a state that has similar S6 helix tilt and is only ∼2.2 Å from the Ca^2+^-bound Cryo-EM structure at the end of the 10 μs run (Figure 2e, red vs. cyan cartoons). These pore structural changes roughly doubled the pore diameter, from ∼10 Å to ∼20 Å at the intracellular entrance (z ∼ − 20 Å; Figure 2f). It should be noted that the full-length BK channel in the Ca^2+^ bound state has an even larger intracellular opening (Figure 2f, green trace), suggesting that additional dilation of the pore may occur at longer timescales, or in response to Ca-binding to the full-length channel. On the other hand, the simulation construct does not include the 11-residue Kv mini-tails required for assembly and membrane insertion of Core-MT (*68*), which could impact the pore conformation. Furthermore, the single-channel conductance of Core-MT is ∼30% lower (*68*), suggesting that its open pore is more constrictive than that of the full-length BK channels.

The opening transitions were accompanied with rehydration of the pore and the channel became conductive to K^+^ (Figure 3 and Movie S3). Consistent with the smaller pore diameter, the final states of the pore in the simulations only accommodate ∼30 waters, compared to ∼40 waters for the Ca^2+^-bound state of full-length BK (*32*). Furthermore, the single-channel conductance estimated from the last 2 μs of *sim 2b* (Figure 3a) is only ∼1.5 pS, much lower than the experimental value of ∼220 pS for Core-MT (*69*). We note that classical force fields, such as CHARMM36m used in the current simulations, are known to underestimate the single channel conductance by about one order of magnitude (*73*). Indeed, the fully opened state of Core-MT, constructed from the Ca^2+^-bound Cryo-EM full-length BK structure, is predicted to have a conductance of ∼6 pS using the same simulation setup (*sim 7*; Figure S3), which is only ∼4 fold of that of the final state from the voltage activation simulations. We further note that subconductance open states have been observed in single channel recordings of BK channels (*74–77*). Besides the limitation of the current fixed charge force fields in quantitively predicting channel conductance, we note that the molecular basis for the large conductance of BK channels is actually poorly understood (*78*). It is noteworthy that the pore hydration level appears to be an important factor in determining the apparent conductance in the simulation, which has also been proposed in a previous atomistic simulation study of the *Aplysia* BK channel (*33*). Despite not reaching a fully conductive state, the ability of the dilated and hydrated pore to sustain K^+^ permeation throughout the second half of the 10 μs simulation demonstrates that the hydrophobic gate in the closed channel has been broken and the channel enters a conductive state. Consistent with previous simulations of other K^+^ channels (*78–80*), the conductance follows a multi-ion mechanism, where there are at least two K^+^ ions inside the filter, preferentially occupying positions S1/S3 or S2/S3 (Figure 3b). The two bound ions subsequently move to S0/S1 when an incoming ion takes the S3 position. Notably, between the ions inside the filter, there can be either one or no water molecule, indicating the coexistence of both soft and hard knock-on mechanisms.

**Figure 3.**
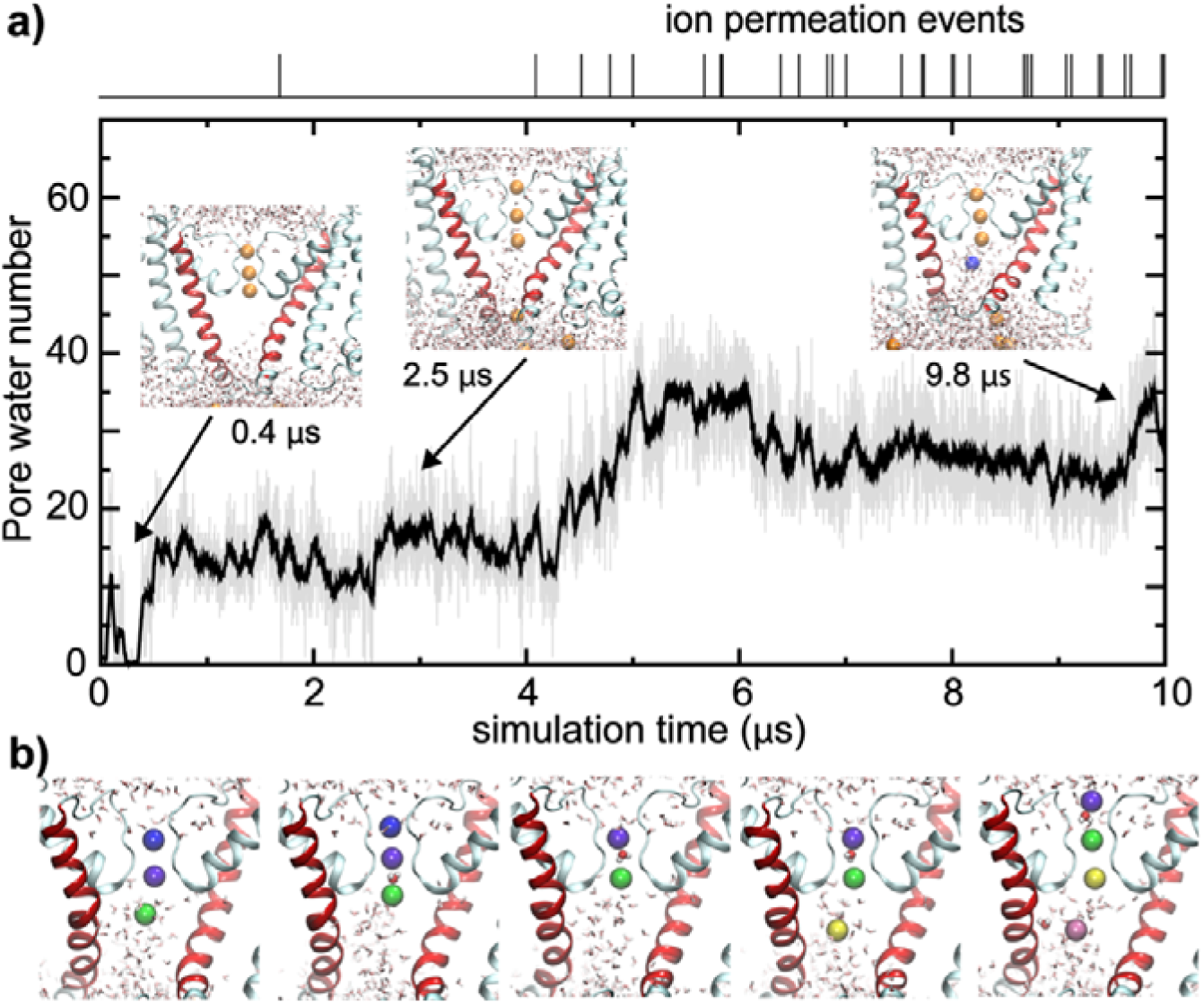
Pore Rehydration and Ion Conductance. **a**) The number of water molecules inside the pore as a function of time during simulation (*sim 2b*), with the upper panel showing recorded ion permeation events. Inserts show snapshots of the pore region at three representative timepoints. **b**) Snapshots illustrating key steps of K^+^ ions passing through the filter. Potassium ions inside or near the filter are colored according to their identities. The water molecule bridging two ions inside the filter is also shown as van der Waals spheres.

### Gating charges: how small VSD movements support voltage sensing

The ability of Anton 2 simulations to directly observe voltage-driven activation of Core-MT BK channels allows one to examine the molecular details of VSD activation and VSD-pore coupling. An important observation is that, even though the charged groups of R210 and R213 show z-displacement of 8-10 Å (Figure 4a), the overall movement of S4 along the membrane normal is only ∼3 Å, less than a single α-helical turn (Figure 2b). This is considerably smaller than that observed in canonical Kv channels, which has been estimated to be ∼ 8 Å (*55*) and up to 15 Å (*56*). Smaller S4 movements have been suggested in fluorophore quenching experiments (*65–67*) and molecular simulations (*39*). Recent Cryo-EM structures of mutant BK channels with constitutively activated VSDs actually reveal minimal S4 movements along the membrane normal (*38, 41*). Upon voltage activation, R210 switches the salt-bridge partner from D186 on S3 to D153 on S2 and D133 on S1, R213 switches to engage with D153, and R207 switches from engaging with D153 and D133 to becoming exposed to the water/membrane interface (Figure 4b). Similar movements were also observed in recent high-resolution Cryo-EM structures of R207A mutant BK channels with constitutively activated VSDs at 0 mV (*41*), even though the net z-displacements of R210 and R213 charges in Cryo-EM structures are about ∼2-4 Å smaller. Both the smaller side chain charge movements and a lack of overall S4 z-displacement in the mutant Cryo-EM structures may be due to the absence of membrane voltage. To further evaluate if z-placements observed at 750 mV are an artifact of unphysical voltage, three independent simulations were initiated from the final state of *sim 2b* at 300 mV for 1.0 μs (*sim 9* in Table S1). The results, summarized in Figure S1, show that the activated state of VSD remained stable and the pore stably hydrated in all three simulations.

**Figure 4.**
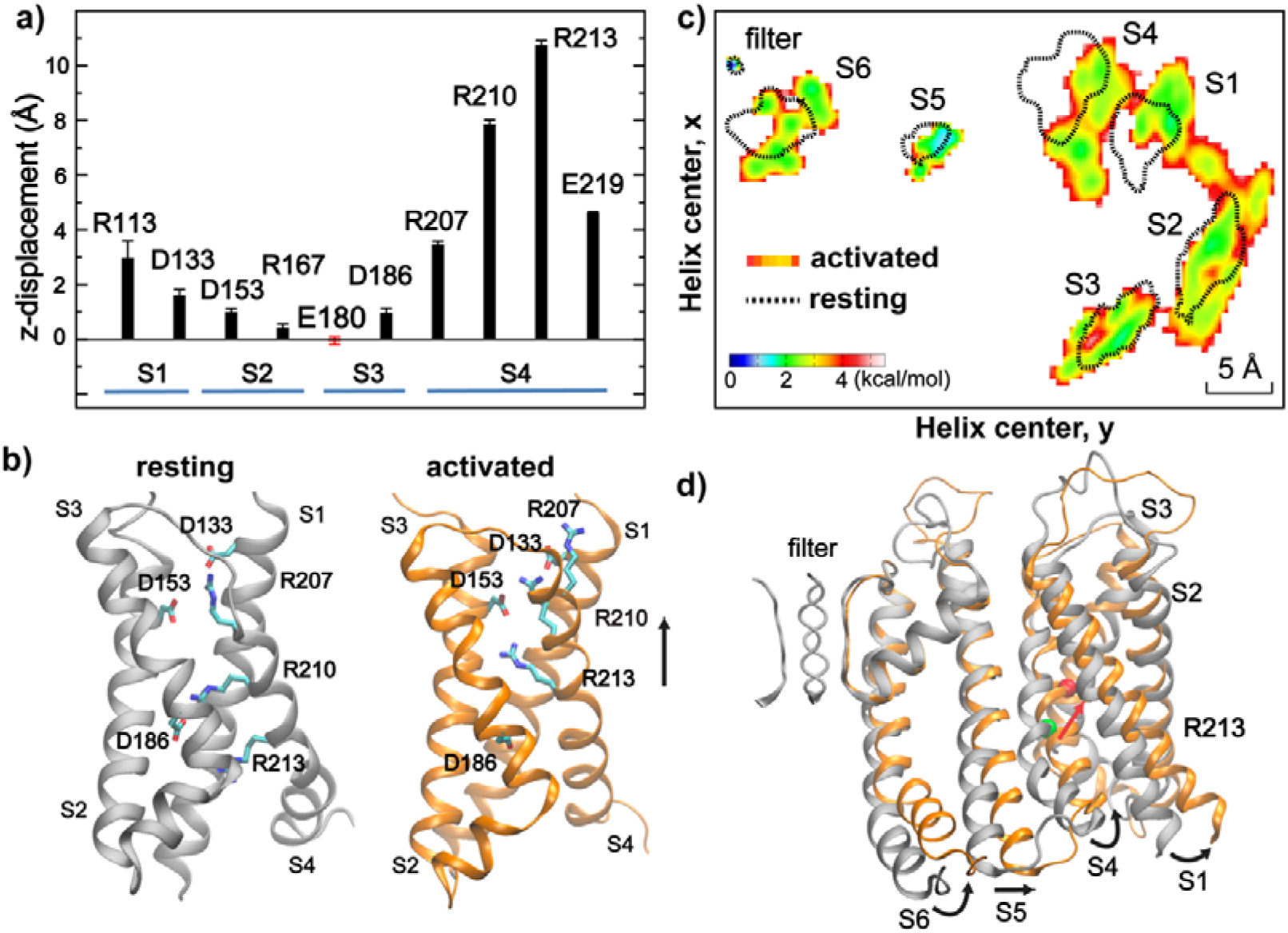
VSD gating charge and voltage-sensing movements. **a)** Average voltage-induced movements of key charges along the membrane normal (z-axis) with respect to the initial resting state structure, derived from last 500 ns of 750 mV simulation *sim 2b*. Error bars show the standard deviations. **b**) Conformations of key charged residues in the resting (silver) and activated (orange) states of BK VSD. The resting and activated states are represented using the snapshots at 0 and 10 _μ_s of sim2b, respectively. **c**) Distributions of centers-of-mass of TM helices along the membrane lateral directions (x and y) (view from the cytosolic side). The distributions for resting and activated states were derived from the first and last 500 ns of the 750 mV simulation *sim 2b*, respectively, which were converted into the free energy scale by ∼ R T ln *P*(*x*,*y*) with T = 300 K. The contour for the resting state distribution (dotted lines) is drawn at 4 kcal/mol. **d**) Overlay of the resting (silver) and activated (orange) states of the TM domain of the Core-MT BK channel. The green and red spheres mark the backbone C_α_ atom of S4 R213 in the resting and activated states. Only one subunit is shown for clarity but all four filter loops shown for reference.

Another important feature is that multiple helices in VSD underwent substantial outward movements in the membrane lateral direction in addition to the modest z-displacement of S4. As shown in Figure 4c, both S1 and S4 displayed ∼3 Å outward movements within the x-y plane. These lateral movements are apparently important for driving the opening motion of the pore-lining S6 helices (Figure 4d), likely mediated by direct S4-S5-S6 interactions (see the next section). Importantly, the predicted VSD movements appear highly consistent with observations from fluorophore quenching experiments (*65–67*), which suggest that S4 likely moves towards S3 and closer to S1/S2 upon activation. Interestingly, the simulations predict that S1 moves during voltage-activation instead of S2, in contrast to the previously proposed model (*66*). Lateral movements of S4 at the cytosolic end have also been observed in recent Cryo-EM structures of R202Q mutant *Aplysia* BK channel (equivalent to R213Q in human BK), which presumably locks the VSDs in the activated state (*38*), even though the overall movements of VSD are much more subtle in the Cryo-EM structures. The later may again be a direct consequence of the absence of membrane voltage.

To understand how the modest movements of VSDs can support effective voltage sensing in BK channels, we calculated the average electrostatic potential maps for the resting and activated states (Figure 5). The results reveal how the protein significantly remodels the local electric field, creating large gradients along both the membrane normal and lateral directions. This is similar the electric field focusing effects proposed for VSDs of both Kv and BK channels (*39, 81*). In particular, residues R210 and R213 reside in a region of high electrostatic potential (> 1100 mV) in the resting state, but move to a region of much lower electrostatic potential (∼ 600 mV) in the activated state. Therefore, despite the modest S4 movements, charges on R210 and R213 can effectively sense a voltage change of ∼500 mV, compared to the total imposed membrane voltage of 750 mV.

**Figure 5.**
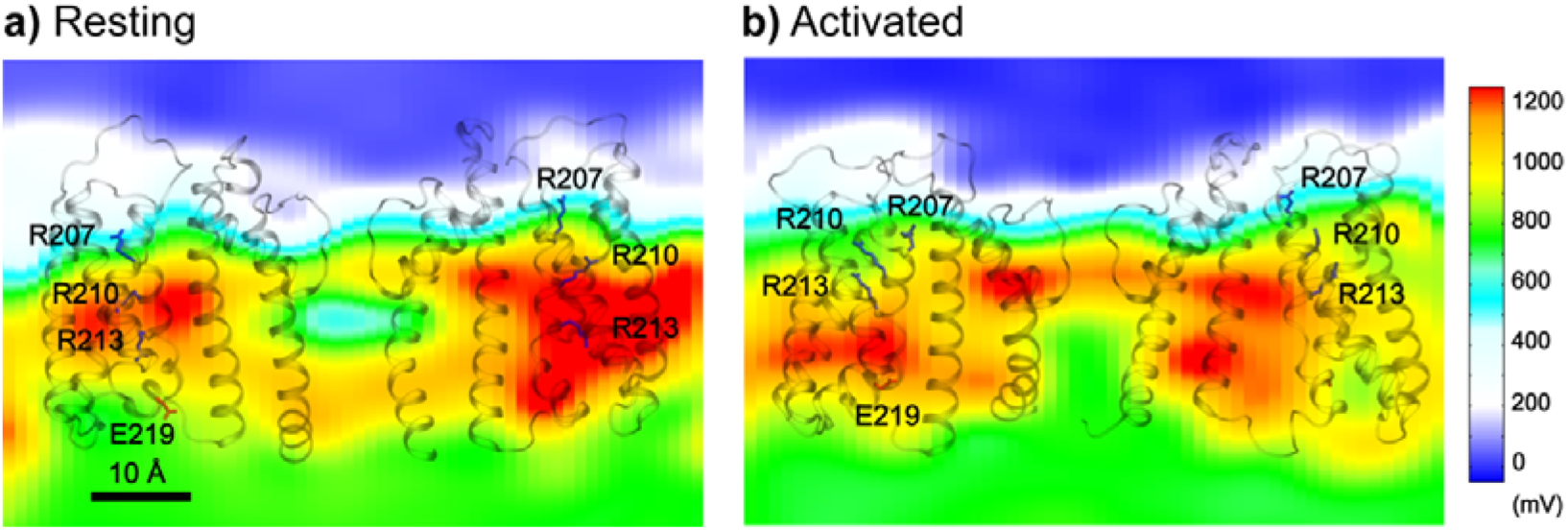
Electrostatic potential fields of the Core-MT BK channel at 750 mV with resting and activated VSDs. The fields were calculated as the averages of the first (resting) and last (activated) 250 ns of simulation *sim2b*. The fields are shown on a plane that goes through the filter and R210. Only two subunits of Core-MT BK are shown for clarity, with side chains of key S4 charges shown in sticks.

We performed free energy calculations (*82, 83*) to further quantify the contributions of key VSD residues to the total gating charge of BK channels (see Methods; Table S1 *sim 3-6*). As summarized in Table 1, the calculations estimated the total gating charge per VSD to be ∼ 0.45 *e*, in strong agreement with the range of 0.48 - 0.65 *e* measured experimentally for the full-length BK channel (*39, 63, 64*). The results further identify R210 and R213 as the primary contributors to voltage sensing, accounting for approximately 97% of the total gating charge per VSD (Table 1). The calculated residue contributions to gating charge are highly consistent with the latest experimental measurements by Carrasquel-Ursulaez et al (2022) (*39*), even though earlier experimental studies disagreed on the contributions of several residues, including R207, R167, D153 and D186. It has been recognized that interpretation of gating charge measurements on mutant BK channels is nontrivial (*39*). For example, neutralizing R213 reduces the total gating charge from 2.62 *e* for WT to 1.31 *e* for R213C; yet replacing it with a negative charge (R213E) does not further reduce the net gating charge (*63*). The simulations reveal that the complication in experimental analysis may be attributed to modest movements of VSD in both membrane normal and lateral directions and a clear interplay of VSD conformation and local electric field (Figure 5). Nonetheless, atomistic simulations together with electrostatic potential and free analyses strongly support a central role of R210 and R213 in BK voltage sensing.

**Table 1.** Residue contributions to the gating charge per VSD.

| Residues | Current work<sup>a</sup> | Carrasquel-Ursulaez (2022)<sup>(39)</sup> | Ma (2006)<sup>(63)</sup> | Diaz (1998)<sup>(64)</sup> |
| --- | --- | --- | --- | --- |
| D153 (S2) | 0.08 ± 0.04 | -0.03 | 0.26 |  |
| R167 (S2) | 0.002 ± 0.03 | -0.02 | 0.15 |  |
| D186 (S3) | -0.01 ± 0.03 | 0.0 | 0.22 |  |
| R207 (S4) | 0.04 ± 0.01 | -0.01 | -0.03 | 0.275 |
| R210 (S4) | 0.25 ± 0.02 | 0.32 to 0.35<sup>b</sup> | -0.01 | 0.325 |
| R213 (S4) | 0.19 ± 0.01 | 0.23 to 0.31<sup>b</sup> | 0.34 | 0.3 |
| E219 (S4) | -0.09 ± 0.02 | 0.03 | 0.02 |  |

<sup>a</sup> Statistical error was determined by block averaging.
<sup>b</sup> Results from reversed charge mutations are excluded here.

The dominant role of R210 and R213 in voltage sensing of BK channels was further tested by steered MD simulations (*sim 8*), where biasing potentials were applied to steer the movement of the guanidium CZ atoms of R210 and R213 from their positions in the initial resting state to those in the activated state in absence of membrane voltage (see Methods and Figure S4). The simulations show that upward movements of R210 and R213 guanidium tips alone can readily drive pore opening, which were observed in two out of four independent steer MD simulations (Figure S4). In the other two simulations, the S5-S6 packing was ruptured during the first 100 ns, likely due to the rapid rate of pulling. Similar to simulations under 750 mV membrane potential, pore dilation and increase in S6 tilting were observed as the VSD responded to the movement of R210 and R213, along with the breakdown of the hydrophobic barrier, as indicated by an increase in the number of water molecules within the pore. The ability of R210 and R213 charge upward movement alone to drive pore opening in absence of membrane voltage further supports the dominant roles of these two charges in voltage sensing and gating of BK channels.

### Central role of the S4-S5-S6 interface in VSD-pore coupling

Examination of the covariance matrices (Figure S5) reveal that all TM helices, particularly, S1-S5, are tightly coupled in both resting and activated states. The lateral movements of S1 and S4 during activation mainly involve the cytosolic ends, concerted with those of S5 and S6 from the pore (Figure 4b). In the Cryo-EM structures, a kink is observed in S4 below R213, resulting in approximately a ∼30° bend in the C-terminal half away from S5 and the pore. During voltage-induced activation, the N-terminal region of S5 maintained strong contacts with the S4 C-terminal segment (Figure S6), such that the upward and outward movement of S4 induces an increase in the tilt and bend of S5 (Fig. 4d). This S4-S5 movement in turn increases the tilt angle of S6 C-terminal half (F315 to E324) by ∼15° to dilate the pore. In particular, the hydrophobic helix-helix contacts between S5 (L235, L239 and F242) and S6 (F315, V319 and I322) are strengthened, while those between N231 and K234 on S5 and E321 and E324 on S6 are weakened during this process (Figure S6). Besides pore dilation, the conformational response of S6 alters the orientation of the conserved E321 and E324 at the cytosolic entrance from membrane interface-facing in the closed state to pore-facing in the open state, as observed in Cryo-EM structures (*28–30*). The resulting increase in both pore size and surface hydrophilicity promotes pore hydration and disrupts vapor barrier to render the channel conductive.

To further identify structural elements central to BK VSD-pore coupling, we performed unbiased dynamic community and coupling pathway analysis based on the covariance matrices derived from atomistic simulations (see Methods). The results, summarized in Figure 6a, reveal that S4-6 forms a single community distinct from the rest of the VSD (and S0 helix), where the motions of residues are more strongly coupled to each other than to the rest of the channel. Indeed, the optimal and suboptimal pathways of allosteric coupling between S4 R213 (a major voltage sensing charge) and S6 E321 (a key position of pore dilation and hydrophilicity increase) pass exclusively through S5 residues (Figure 6b). The central role of S5 in mediating VSD-pore coupling is further supported by analyzing the information flow betweenness (*84*), which provides a global measures how conformational perturbation flows through the network from the “source” (*e.g.,* R213) to the “sink” (e.g., E321). The analysis reveals that S5 residues have the largest contributions to the informational flow besides neighboring residues on S4 or S6 (Figure 6c). Other residues on S1 or S3 also contribute significantly to the flow, albeit at much lower levels compared to S5. Interestingly, analysis of the simulation trajectory of Ca^2+^ fully open state (Table S1, *sim 7*) reveals highly similar patterns in the dynamic coupling communities, coupling pathways as well as informational flow (Figure S7), despite substantial differences in the VSD conformation and S4-S5-S6 packing (Figure S8). The implication is that Ca²_⁺_- and voltage activation pathways likely have significant overlaps, which is consistent with experimental observations that BK channels can be synergistically or independently activated by membrane depolarization and Ca²_⁺_ binding. Taken together, the simulation and dynamic network analysis consistently point to a central role of the S4-S5-S6 interface in VSD-pore coupling of BK channels.

**Figure 6:**
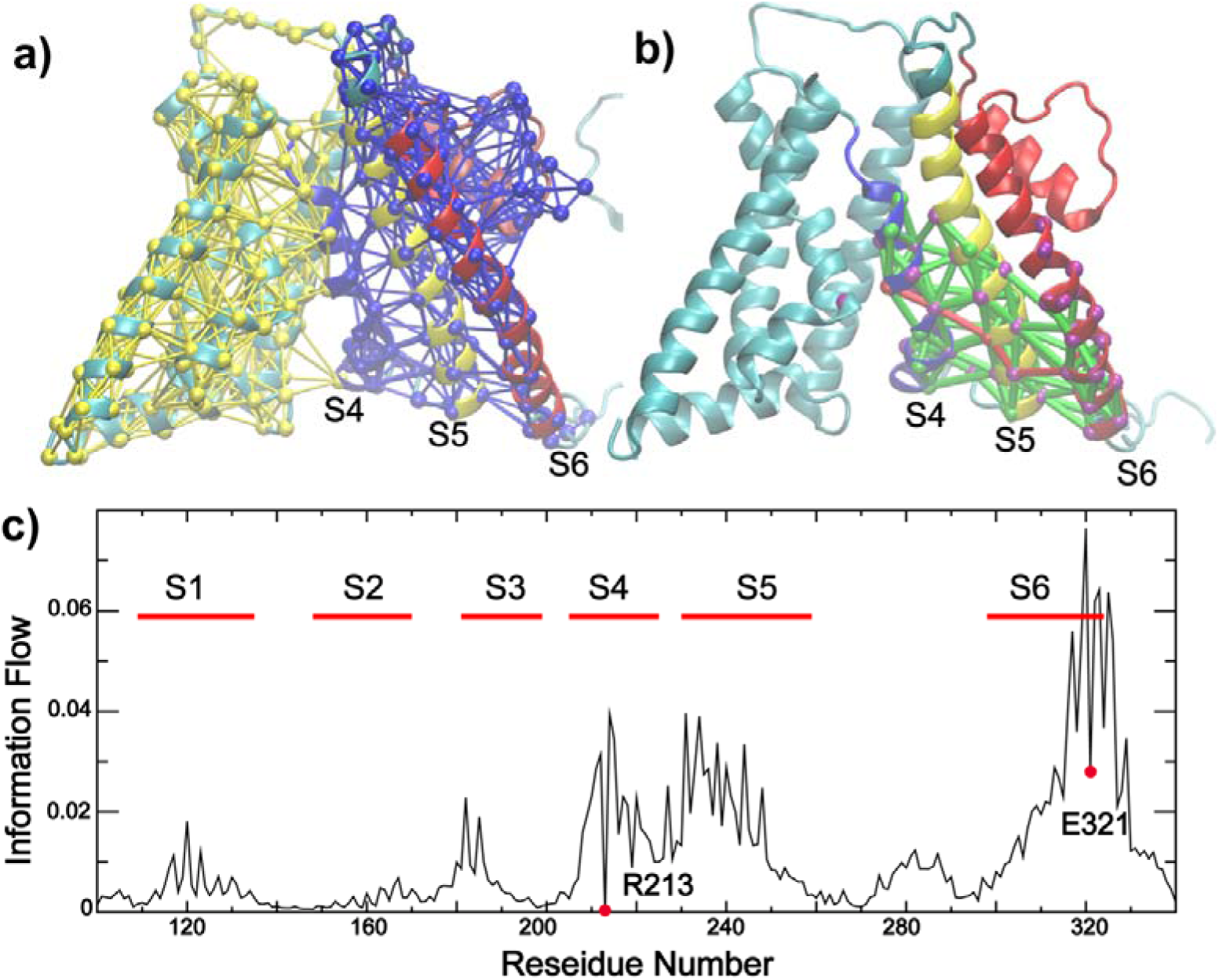
Dynamic community, coupling pathways, and information flow of VSD-pore coupling in BK. **a)** Dynamic community analysis showing that TM S4-6 are clustered into single tightly coupled community (blue network). The nodes (residues) and edges (contacts) are colored based on the community number. **b)** Optimal and suboptimal pathways of dynamic coupling between R213 (VSD S4) and E321 (pore-lining S6). All paths are colored green except for the optimal path, which is colored red. Nodes with information flow value > 0.02 is highlighted in purple; **c)** Information flow profile of the Core-MT BK channel with R213 as the source and E321 as the sink node (labeled by red circle), respectively. All dynamic coupling analysis was derived from last 500 ns of *sim 1* (closed state at 0 mV; see Table S1).

## Concluding Discussion

Long atomistic simulations in explicit water and membrane performed using special purposed supercomputer Anton 2 have allowed direct observation of voltage-induced activation of the Core-MT BK channels for the first time. These simulations provide crucial new insights into how BK VSDs may sense membrane voltage and how the VSD movements may drive the pore opening and activate channel conductance. In particular, the simulations support that the S4 helix remains the major voltage sensing element and that it undergoes modest movements of ∼ 3 Å along both the membrane normal and lateral directions. The ability of BK channels to utilize relatively small VSD movements to sense membrane voltage and drive channel activation is likely attributed to the protein’s ability to modify the local electric field, which leads to large electrostatic potential gradient in both vertical and lateral directions. Further analyses reveal that R210 and R213 are the major voltage sensing residues in the wild-type Core-MT BK channel, contributing ∼97% to the total gating charge of ∼ 0.45 *e* per VSD. These core features of BK voltage sensing from simulation are largely consistent with a range of existing functional, biophysical and structural studies (*38, 39, 41, 66*), even though structural studies mutant BK channels with constitutively activated VSDs reveal smaller movements of R210 and R213 guanidinium moieties and minimal vertical movement of S4. The later observations may be a consequence of the complete absence of membrane voltage during structural determination. Importantly, control simulations show that the activated VSD state remains stable at 300 mV, suggesting that the observed vertical S4 movement is not an artifact of high voltage of 750 mV.

Another distinct feature derived from atomistic simulation is that the tightly packed S4-S5-S6 interface in BK channels plays a central role in the VSD-pore coupling, which is further supported by dynamic community, coupling pathway, and informational flow analyses. The modest movements in S4 drive a series of concerted conformational changes in S5 and eventually in S6, with the S4-S5 linker not playing a major role as observed in the canonical Kv channels (*55–57*). This is supported by the findings that mutations in the S4-S5 linker (N225-K228) do not significantly affect the coupling between VSD and the pore (*40*). Arguably, the distinct features of voltage-sensing and pore-sensor coupling observed in BK channels are intimately related to the novel non-domain-swapped TM topology. In particular, the VSD and pore domains are more tightly packed compared to the domain swapped configuration (Figure 1), which restricts the VSD motions and provides a direct pathway for their inter-talk. Interestingly, a similar non-canonical pathway for VSD-pore coupling, involving the S4/S1 and S1/S5 interfaces, has recently been proposed for the non-domain-swapped cardiac hERG potassium channel (*85*). We also note that a similar non-canonical VSD-pore coupling involving S4-S5 interactions between neighboring subunits has been suggested to complement the canonical coupling mode even in domain-swapped Kv channels (*57, 86, 87*). An increasing number of ion channels have been discovered to adopt non-domain-swapped TM topology besides BK, including hERG (*88*), KvAP (*89*), HCN (*90*), Eag1 (*91*), and CNG channels (*92*). It is possible that the mechanistic features observed for BK channels may apply to voltage gating of the important emerging class of non-domain swapped ion channels in general.

Importantly, full-length BK channels can be independently activated by membrane potential and intracellular Ca^2+^ (*22, 93, 94*). The latter requires the C-terminal gating ring that is absent in the Core-MT construct studied in this work. Cryo-EM structures have revealed functional state-dependent interactions between the gating ring and VSD (*28–30, 38*), strongly supporting that the calcium sensor and VSD are coupled. It is not clear how the VSD-gating ring coupling may affect the nature of VSD movements or how VSDs may drive the pore opening. Nonetheless, mechanistic details revealed from the voltage-gating of the Core-MT construct should provide a solid basis for future computational and experimental studies of BK activation and regulation.

## Methods and Materials

### Molecular modeling and atomistic simulations

The structure of the Core-MT human BK channel in the closed state was derived from the Cryo-EM structures of the *ac*BK channel in deactivated Ca^2+^-free states (PDB 5tji (*29*)) as previously described (*32, 95*). The simulated construct was truncated at R342. The 11-residue C-terminal Kv mini-tails are not involved in voltage gating (*68, 69*), and they were thus not included. The dynamic loop (C54-V91) and N-terminal tail (M1-N19) were not included either. Residues before and after the missing segments are capped with either an acetyl group (for N-terminus) or a N-methyl amide (for C-terminus). Standard protonation states under neutral pH were assigned for all titratable residues.

The initial structures was first inserted in model POPC lipid bilayers and then solvated in TIP3P water using the CHARMM-GUI web server (*96*). Even though polarizable water models are probably necessary to capture the precise energetics (and kinetics) of dewetting transitions within a protein cavity, it has also been shown that classical nonpolarizable water models such as TIP3P is sufficient to capture the spontaneous dewetting and rehydration of BK channels (*32–34, 36*). In fact, a recent analysis using the Drude polarizable force field actually under-estimated the hydration free energy of deactivated BK pore (∼0.5 kcal/mol) (*35*), which is very likely too small to sustain a dry and nonconductive pore. All systems were neutralized and 150 mM KCl added. The final simulation boxes contain about 593 lipid molecules (POPC) and ∼50,000 water molecules and other solutes, with a total of ∼250,000 atoms and dimensions of ∼160 × 160 × 110 Å^3^. The CHARMM36m all-atom force field (*97*) and the CHARMM36 lipid force field (*98*) were used. All simulations were performed using Desmond (*99*) on Anton 2 (*70, 71*) or CUDA-enabled versions of Gromacs 2020 (*100, 101*) on GPU clusters. Electrostatic interactions were described by using the Particle Mesh Ewald (PME) algorithm (*102*) with a cutoff of 12 Å. Van der Waals interactions were cutoff at 12 Å with a smooth switching function starting at 10 Å. Covalent bonds to hydrogen atoms were constrained by the SHAKE algorithm (*103*), and the MD time step was set at 2 fs. The temperature was maintained at 298 K using the Nose-Hoover thermostat (*104, 105*) (in Gromacs). The pressure was maintained semi-isotopically at 1 bar at membrane lateral directions using the Parrinello–Rahman barostat algorithm (*106*).

All systems were first minimized for 5000 steps using the steepest descent algorithm, followed by a series of equilibration steps where the positions of heavy atoms of the protein and lipid were harmonically restrained as prescribed by the CHARMM-GUI Membrane Builder (*107*). For Anton simulation, additional 70 ns equilibration step with 0.1 kcal.mol^-1^.Å^-2^ position restraints on all protein heavy atoms was performed before production runs. Note that the inner pore became dewetted during equilibration simulations. All production simulations were performed under NVT (constant particle number, volume and temperature) conditions at 298 K. The P-loop/filter (T273 to D292) and C-terminus were harmonically restrained with a small force constant of 0.1 kcal.mol^-1^.Å^-2^ in all production simulations to prevent drift in all production simulations. Using Anton 2, a 2-μs control simulation was first performed without membrane voltage (*sim 1*, Table S1). Two independent 10-μs simulations were then performed at 750 mV to directly probe voltage-driven activation transitions (*sim 2*). The voltage was applied as a constant external electric field (*E* = *V*/*L*_z_) imposed along the z dimension (*83*).

### Steered molecular dynamics simulations

Steered molecular dynamics (SMD) simulations were performed to test the role of R210 and R213 in voltage sensing and channel activation. In these simulations, a moving reference point was used to simulate the effect of an external electric field by pulling the CZ atoms of R210 and R213 from their initial positions in the resting state to those observed in the activated state. As illustrated in Figure S4, the initial reference points of SMD were positioned at the location of each CZ atom of R210 and R213 in the resting/closed state. From 0 to 100 ns, these reference points were moved along the z-axis at constant speed until they reached the positions corresponding to the predicted activated VSD state at 100 ns. A harmonic positional restraint of 5 kcal/mol·Å² was applied between each CZ atom to its respective reference point along z-axis, allowing the CZ atom to track the reference point’s movement. After 100 ns, the reference points were held stationary at the positions corresponding to the activated VSD state, with the same harmonic restraints imposed to maintain the CZ atoms in their activated position. Four independent simulations (*sim 8*) were performed, and pore opening was observed in two of the four replicates (replicas 2 and 4). In the other two replicas, the S5-S6 packing was broken during the first 100 ns, likely due to strong steering potentials imposed, and the simulations were terminated.

### Free energy analysis of gating charges

The total gating charge ΔQ is the sum of contributions of all charged residues in VSD and can be calculated as (*82, 83*):

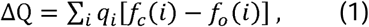

where is *q_i_* is the charge of a specific residue, and the dimensionless quantities *f_s_(i)* for a specific conformational state *s*, which can be open (*o*) or closed (*c*), represent the coupling of charge *q_i_* to the transmembrane potential. The value of *f_s_(i)* is derived from the difference between charging free energies calculated at two different voltages *V*_1_ and *V*2,

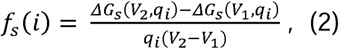

where *ΔG_s_(V, q_i_ )* is the free energy cost of increasing the charge of residue *i* from 0 to *q_i_* in state *s* under membrane voltage *V. ΔG_s_(V, q_i_ )* was calculated using thermodynamic integration (TI), with the charge gradually scaled to the final value in increments of 0.1, *i.e.*, λ = 0, 0.1, 0.2, …., 1.0. In this work, we focused on 7 charged residues on VSD, namely, D153, R167, D186, R207, R210, R213 and E219, and performed four TI free energy calculations for each residue (sim 3-6, Table S1). The last snapshot of simulation 2b was used to represent the activated state of VSD. For each TI window, the system was first equilibrated 200 ps, and then simulated for 4 ns to calculate the mean force <δH/δλ>_λ_.

### Structural and electrostatic potential analysis

The displacement of residue charged groups, as well as movement of helixes (translocation, titling and rotation) is calculated using MDanalysis (*108*) together with in-house scripts. The TM helices are defined as: T109-S135 (S1), F148-A170 (S2), V181-L199 (S3), G205-N225 (S4), S230-S259 (S5), and A313-E324 (S6 after the glycine hinge). Residue contacts were identified using a minimal heavy atom distance cutoff of 5 Å. Pore water molecules were identified as those occupying the inner pore cavity below the selectivity filter, roughly from L312 to the plane defined by the center of mass of P320. Pore profiles are calculated using program HOLE(*109*). To examine how the protein modulates the local electric field, average electrostatic potential maps were then calculated from the first (resting) and last (activated) 250 ns of *sim 2b* using a combination of Python scripts and the VMD’s PMEPot and VolMap plugins (*110*). The electrostatic potential is obtained by solving the Poisson equation 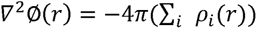 where the sum runs over all atoms, and ρ*_i_(r)* is the charge electric potential distribution contributed by atom *i* at position *r* approximated by a spherical Gaussian: 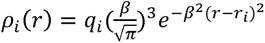. An Ewald factor β of 0.25 Å^-1^ was used for the inverse width of the Gaussian and grid size was set to 1 Å. All molecular illustrations were prepared using VMD (*111*). External electric potential was added to the PMEPot grid file with in-house scripts (see supplementary files).

### Dynamic community and coupling pathways

Dynamic network analysis was performed using the *Networkview* (*112*) plugin of VMD . To build the network, each amino acid was represented as a single node at the Cα position, and a contact (edge) was defined between two nodes if the minimal heavy-atom distance between residues was within a cutoff distance (5 Å) during at least 75% of the trajectory. The resulting contact matrix was weighted based on the covariance of dynamic fluctuation (*C_ij_*) calculated from the same MD trajectory as *w_ij_* = - log(|*C_ij_*|). The length of a possible pathway *D_ij_* between distant nodes *i* and *j* is defined as the sum of the edge weights between consecutive nodes along this path.

The shortest path, calculated using Floyd-Warshall algorithm (*113*), is considered the optimal pathway with the strongest dynamic coupling. Suboptimal paths are identified as alternative top-ranked paths with lengths that deviate by less than 50% from the optimal path. We further performed dynamic community analysis to identify groups of residues are more tightly coupled among themselves (*114*).

### Information flow analysis

Information flow provides a global assessment of the contributions of all nodes to the dynamic coupling between selected “source” and “sink” nodes(*115, 116*). This analysis complements the dynamic pathway analysis to provide additional insights on how different residues may contribute to sensor-pore coupling. For this, a network similar to the one described above was first constructed. Pairwise mutual information was calculated between node *i* and *j* as follows: *M_ij_ = H_i_ + H_j_ − H_ij_*. *H_i_* is calculated as 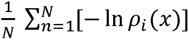, where ρ*_i_(x)* is the fluctuation density and *x* is the distance to the equilibrium position(*116*). Gaussian mixture model (GMM)(*^117^*) is used to estimate the density. The residue network is then defined as *A*_ij_ = *C*_ij_ *M*_ij_, where *C_ij_* is the contact map. To analysis the information flow from the source (*S_0_*) to sink (*S_I_*) nodes, the network Laplacian, is calculated as *L* = *D* - *A*, where D is diagonal degree matrix: *D_ii_* = ∑*_J_ A_iJ_* . The information flow through a given node (residue) is defined as 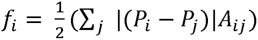. The potentials *P* is given by P = *L*^∼-1^ b, where *L*^∼-1^ is the inverse reduced Laplacian, and *b* is the supply vector that corresponds to one unit of current entering at the source node that will exit at sink nodes. The magnitude of *f_i_* thus quantifies the contribution of residue *i* to dynamic coupling between the source and sink nodes.

## Acknowledgements

The authors thank Jianmin Cui and Guohui Zhang for critical discussions. This work is supported by NIH R35 GM144045 (Chen). Anton 2 computer time was provided by the Pittsburgh Supercomputing Center (PSC) through Grant R01 GM116961 from the National Institutes of Health. The Anton 2 machine at PSC was generously made available by D.E. Shaw Research.

## Author contributions

Chen and Jia conceived and initiated the study. Jia performed simulations and analysis. Jia and Chen wrote the manuscript.

## Competing interests

The authors declare no competing interests.

## Data availability statement

All data needed to evaluate the conclusions in the paper are present in the paper and the Supplementary Materials. The PDB structures of Core-MT with resting and activated VSDs, as well as simulation input and analysis script, can be found on GitHub at: https://github.com/mdlab-um/Votage_gating_Core-MT.

## Supplementary information

The online version contains supplementary material available at https://doi.org/xxxx

## Notes

### Competing Interest Statement

The authors have declared no competing interest.

### Summary of Updates

Figure S8 updated to correct residue labels

## References

1. A. L. Hodgkin, A. F. Huxley, A quantitative description of membrane current and its application to conduction and excitation in nerve. J. Physiol. 117, 500–544 (1952).

2. F. Bezanilla, The voltage sensor in voltage-dependent ion channels. Physiol. Rev. 80, 555–592 (2000).

3. F. J. Sigworth, Structural biology: Life’s transistors. Nature 423, 21–22 (2003).

4. R. MacKinnon, Potassium channels. FEBS Lett. 555, 62–65 (2003).

5. K. J. Swartz, Sensing voltage across lipid membranes. Nature 456, 891–897 (2008).

6. J. E. Brayden, M. T. Nelson, Regulation of arterial tone by activation of calcium-dependent potassium channels. Science 256, 532–535 (1992).

7. G. C. Wellman, M. T. Nelson, Signaling between SR and plasmalemma in smooth muscle: sparks and the activation of Ca^2+^-sensitive ion channels. Cell Calcium 34, 211–229 (2003).

8. S. Pluger et al., Mice with disrupted BK channel beta1 subunit gene feature abnormal Ca^2+^ spark/STOC coupling and elevated blood pressure. Circ Res 87, E53–60. (2000).

9. M. T. Nelson, J. M. Quayle, Physiological roles and properties of potassium channels in arterial smooth muscle. Am. J. Physiol. 268, C799–822 (1995).

10. Y. Tanaka, M. Aida, H. Tanaka, K. Shigenobu, L. Toro, Involvement of maxi-K_Ca_ channel activation in atrial natriuretic peptide-induced vasorelaxation. Naunyn Schmiedebergs Arch Pharmacol 357, 705–708 (1998).

11. G. J. Perez, A. D. Bonev, J. B. Patlak, M. T. Nelson, Functional coupling of ryanodine receptors to K_Ca_ channels in smooth muscle cells from rat cerebral arteries. J Gen Physiol 113, 229–238 (1999).

12. S. Sokolov, T. Scheuer, W. A. Catterall, Gating pore current in an inherited ion channelopathy. Nature 446, 76–78 (2007).

13. M. J. Ackerman, The long QT syndrome: Ion channel diseases of the heart. Mayo Clin. Proc. 73, 250–269 (1998).

14. G. J. Kaczorowski, M. L. Garcia, Pharmacology of voltage-gated and calcium-activated potassium channels. Curr. Opin. Chem. Biol. 3, 448–458 (1999).

15. F. Lehmann-Horn, K. Jurkat-Rott, Voltage-gated ion channels and hereditary disease. Physiol. Rev. 79, 1317–1372 (1999).

16. S. C. Cannon, in Annu. Rev. Neurosci. (2006), vol. 29, pp. 387–415.

17. B. S. Meldrum, M. A. Rogawski, Molecular targets for antiepileptic drug development. Neurotherapeutics 4, 18–61 (2007).

18. C. A. Reid, S. F. Berkovic, S. Petrou, Mechanisms of human inherited epilepsies. Prog. Neurobiol. 87, 41–57 (2009).

19. N. Eijkelkamp et al., Neurological perspectives on voltage-gated sodium channels. Brain 135, 2585–2612 (2012).

20. D. L. H. Bennett, C. G. Woods, Painful and painless channelopathies. Lancet Neurol. 13, 587–599 (2014).

21. A. Marty, Ca-dependent K channels with large unitary conductance in chromaffin cell membranes. Nature 291, 497–500 (1981).

22. F. T. Horrigan, R. W. Aldrich, Coupling between Voltage Sensor Activation, Ca2+ Binding and Channel Opening in Large Conductance (BK) Potassium Channels. The Journal of General Physiology 120, 267–305 (2002).

23. K. L. Magleby, Gating mechanism of BK (Slo1) channels: So near, yet so far. The Journal of General Physiology 121, 81–96 (2003).

24. L. Salkoff, A. Butler, G. Ferreira, C. Santi, A. Wei, High-conductance potassium channels of the SLO family. Nat Rev Neurosci 7, 921–931 (2006).

25. G. F. Contreras et al., A BK (Slo1) channel journey from molecule to physiology. Channels 7, 442–458 (2013).

26. B. H. Bentzen, S.-P. Olesen, L. C. B. Rønn, M. Grunnet, BK channel activators and their therapeutic perspectives. 5, (2014).

27. H. Yang, G. Zhang, J. Cui, BK channels: multiple sensors, one activation gate. Front Physiol 6, 29 (2015).

28. R. K. Hite, X. Tao, R. MacKinnon, Structural basis for gating the high-conductance Ca2+-activated K+ channel. Nature 541, 52–57 (2017).

29. X. Tao, R. K. Hite, R. MacKinnon, Cryo-EM structure of the open high-conductance Ca2+-activated K+ channel. Nature 541, 46–51 (2017).

30. X. Tao, R. MacKinnon, Molecular structures of the human Slo1 K(+) channel in complex with beta4. Elife 8, (2019).

31. S. B. Long, E. B. Campbell, R. Mackinnon, Crystal structure of a mammalian voltage-dependent Shaker family K+ channel. Science 309, 897–903 (2005).

32. Z. Jia, M. Yazdani, G. Zhang, J. Cui, J. Chen, Hydrophobic gating in BK channels. Nat. Commun. 9, 3408 (2018).

33. R. X. Gu, B. L. de Groot, Central cavity dehydration as a gating mechanism of potassium channels. Nat. Commun. 14, (2023).

34. E. B. Nordquist, Z. Jia, J. Chen, Inner pore hydration free energy controls the activation of big potassium channels. Biophys. J. 122, 1158–1167 (2023).

35. J. Deng, Q. Cui, Electronic Polarization Leads to a Drier Dewetted State for Hydrophobic Gating in the Big Potassium Channel. The journal of physical chemistry letters, 7436–7441 (2024).

36. L. Coronel, G. D. Muccio, B. Rothberg, A. Giacomello, V. Carnevale, Lipid-mediated hydrophobic gating in the BK potassium channel. ArXiv, (2024).

37. Y. Zhou, H. Yang, J. Cui, C. J. Lingle, Threading the biophysics of mammalian Slo1 channels onto structures of an invertebrate Slo1 channel. J. Gen. Physiol. 149, 985–1007 (2017).

38. F. C. Gustavo, S. Rong, L. Ramon, P. Eduardo, Structural Basis of Voltage-Dependent Gating in BK Channels and Its Coupling to the Calcium Sensor. bioRxiv, 2023.2012.2029.573674 (2023).

39. W. Carrasquel-Ursulaez et al., Mechanism of voltage sensing in Ca(2+)- and voltage-activated K(+) (BK) channels. Proc. Natl. Acad. Sci. U. S. A. 119, e2204620119 (2022).

40. L. Sun, F. T. Horrigan, A gating lever and molecular logic gate that couple voltage and calcium sensor activation to opening in BK potassium channels. Sci Adv 8, eabq5772 (2022).

41. G. S. Kallure, K. Pal, Y. Zhou, C. J. Lingle, S. Chowdhury, High-resolution structures illuminate key principles underlying voltage and LRRC26 regulation of Slo1 channels. bioRxiv, (2023).

42. H. P. Larsson, O. S. Baker, D. S. Dhillon, E. Y. Isacoff, Transmembrane movement of the Shaker K+ channel S4. Neuron 16, 387–397 (1996).

43. O. Yifrach, R. MacKinnon, Energetics of pore opening in a voltage-gated K+ channel. Cell 111, 231–239 (2002).

44. D. Schmidt, Q.-X. Jiang, R. MacKinnon, Phospholipids and the origin of cationic gating charges in voltage sensors. Nature 444, 775–779 (2006).

45. L. D. Islas, F. J. Sigworth, Voltage sensitivity and gating charge in Shaker and Shab family potassium channels. J. Gen. Physiol. 114, 723–741 (1999).

46. W. A. Catterall, Ion channel voltage sensors: structure, function, and pathophysiology. Neuron 67, 915–928 (2010).

47. W. A. Catterall, N. Zheng, Deciphering voltage-gated Na(+) and Ca(2+) channels by studying prokaryotic ancestors. Trends Biochem. Sci. 40, 526–534 (2015).

48. J. Wu et al., Structure of the voltage-gated calcium channel Ca(v)1.1 at 3.6 A resolution. Nature 537, 191–196 (2016).

49. M. J. Lenaeus et al., Structures of closed and open states of a voltage-gated sodium channel. Proc. Natl. Acad. Sci. U. S. A. 114, E3051–E3060 (2017).

50. D. Matthies et al., Single-particle cryo-EM structure of a voltage-activated potassium channel in lipid nanodiscs. Elife 7, (2018).

51. N. E. Schoppa, K. Mccormack, M. A. Tanouye, F. J. Sigworth, The Size of Gating Charge in Wild-Type and Mutant Shaker Potassium Channels. Science 255, 1712–1715 (1992).

52. S. K. Aggarwal, R. MacKinnon, Contribution of the S4 segment to gating charge in the Shaker K+ channel. Neuron 16, 1169–1177 (1996).

53. S. B. Long, X. Tao, E. B. Campbell, R. MacKinnon, Atomic structure of a voltage-dependent K+ channel in a lipid membrane-like environment. Nature 450, 376–382 (2007).

54. S. A. Seoh, D. Sigg, D. M. Papazian, F. Bezanilla, Voltage-sensing residues in the S2 and S4 segments of the Shaker K+ channel. Neuron 16, 1159–1167 (1996).

55. P. W. Fowler, M. S. P. Sansom, The pore of voltage-gated potassium ion channels is strained when closed. Nat. Commun. 4, 1872 | DOI: 1810.1038/ncomms2858 (2013).

56. M. O. Jensen et al., Mechanism of voltage gating in potassium channels. Science 336, 229–233 (2012).

57. P. Hou et al., Two-stage electro-mechanical coupling of a KV channel in voltage-dependent activation. Nat. Commun. 11, 676 (2020).

58. F. T. Horrigan, R. W. Aldrich, Allosteric voltage gating of potassium channels II. Mslo channel gating charge movement in the absence of Ca(2+). J. Gen. Physiol. 114, 305–336 (1999).

59. Y. Lorenzo-Ceballos, W. Carrasquel-Ursulaez, K. Castillo, O. Alvarez, R. Latorre, Calcium-driven regulation of voltage-sensing domains in BK channels. Elife 8, e 44934 (2019).

60. W. Carrasquel-Ursulaez et al., Hydrophobic interaction between contiguous residues in the S6 transmembrane segment acts as a stimuli integration node in the BK channel. J. Gen. Physiol. 145, 61–74 (2015).

61. G. F. Contreras, A. Neely, O. Alvarez, C. Gonzalez, R. Latorre, Modulation of BK channel voltage gating by different auxiliary β subunits. Proceedings of the National Academy of Sciences 109, 18991–18996 (2012).

62. E. Stefani et al., Voltage-controlled gating in a large conductance Ca2+-sensitive K+channel (hslo). Proc. Natl. Acad. Sci. U. S. A. 94, 5427–5431 (1997).

63. Z. Ma, X. J. Lou, F. T. Horrigan, Role of charged residues in the S1-S4 voltage sensor of BK channels. J. Gen. Physiol. 127, 309–328 (2006).

64. L. Diaz et al., Role of the S4 segment in a voltage-dependent calcium-sensitive potassium (hSlo) channel. J. Biol. Chem. 273, 32430–32436 (1998).

65. A. Pantazis, A. P. Kohanteb, R. Olcese, Relative motion of transmembrane segments S0 and S4 during voltage sensor activation in the human BK(Ca) channel. J. Gen. Physiol. 136, 645–657 (2010).

66. A. Pantazis, R. Olcese, Relative transmembrane segment rearrangements during BK channel activation resolved by structurally assigned fluorophore-quencher pairing. J. Gen. Physiol. 140, 207–218 (2012).

67. N. Savalli, A. Kondratiev, L. Toro, R. Olcese, Voltage-dependent conformational changes in human Ca(2+)- and voltage-activated K(+) channel, revealed by voltage-clamp fluorometry. Proc. Natl. Acad. Sci. U. S. A. 103, 12619–12624 (2006).

68. G. Budelli, Y. Y. Geng, A. Butler, K. L. Magleby, L. Salkoff, Properties of Slo1 K+ channels with and without the gating ring. Proc. Natl. Acad. Sci. U. S. A. 110, 16657–16662 (2013).

69. G. Zhang et al., Deletion of cytosolic gating ring decreases gate and voltage sensor coupling in BK channels. J. Gen. Physiol. 149, 373–387 (2017).

70. D. E. Shaw et al., Anton, a special-purpose machine for molecular dynamics simulation. Commun. ACM 51, 91–97 (2008).

71. D. E. Shaw et al., Anton 2: Raising the bar for performance and programmability in a special-purpose molecular dynamics supercomputer. Int Conf High Perfor, 41–53 (2014).

72. T. Jiang, K. Yu, H. C. Hartzell, E. Tajkhorshid, Lipids and ions traverse the membrane by the same physical pathway in the nhTMEM16 scramblase. Elife 6, e28671 (2017).

73. S. Furini, C. Domene, Critical Assessment of Common Force Fields for Molecular Dynamics Simulations of Potassium Channels. J. Chem. Theory Comput. 16, 7148–7159 (2020).

74. W. B. Ferguson, O. B. McManus, K. L. Magleby, Opening and closing transitions for BK channels often occur in two steps via sojourns through a brief lifetime subconductance state. Biophys. J. 65, 702–714 (1993).

75. X. P. Sun, L. C. Schlichter, E. F. Stanley, Single-channel properties of BK-type calcium-activated potassium channels at a cholinergic presynaptic nerve terminal. Journal of Physiology-London 518, 639–651 (1999).

76. V. Gonzalez-Perez, X. H. Zeng, K. Henzler-Wildman, C. J. Lingle, Stereospecific binding of a disordered peptide segment mediates BK channel inactivation. Nature 485, 133–136 (2012).

77. Z. Takacs, J. P. Imredy, J. P. Bingham, B. S. Zhorov, E. G. Moczydlowski, Interaction of the BKCa channel gating ring with dendrotoxins. Channels (Austin) 8, 421–432 (2014).

78. A. Mironenko, U. Zachariae, B. L. de Groot, W. Kopec, The Persistent Question of Potassium Channel Permeation Mechanisms. J. Mol. Biol. 433, 167002 (2021).

79. S. Bernèche, B. Roux, Energetics of ion conduction through the K+ channel. Nature 414, 73–77 (2001).

80. B. Roux, Ion conduction and selectivity in K+ channels. Annu. Rev. Biophys. Biomol. Struct. 34, 153–171 (2005).

81. D. M. Starace, F. Bezanilla, A proton pore in a potassium channel voltage sensor reveals a focused electric field. Nature 427, 548–553 (2004).

82. F. Khalili-Araghi et al., Calculation of the gating charge for the Kv1.2 voltage-activated potassium channel. Biophysical Journal 98, 2189–2198 (2010).

83. B. Roux, The membrane potential and its representation by a constant electric field in computer simulations. Biophys. J. 95, 4205–4216 (2008).

84. W. M. Botello-Smith, Y. Luo, Robust Determination of Protein Allosteric Signaling Pathways. J. Chem. Theory Comput. 15, 2116–2126 (2019).

85. C. A. Z. Bassetto, F. Costa, C. Guardiani, F. Bezanilla, A. Giacomello, Noncanonical electromechanical coupling paths in cardiac hERG potassium channel. Nature Communications 14, 1110 (2023).

86. A. I. Fernandez-Marino, T. J. Harpole, K. Oelstrom, L. Delemotte, B. Chanda, Gating interaction maps reveal a noncanonical electromechanical coupling mode in the Shaker K(+) channel. Nat Struct Mol Biol 25, 320–326 (2018).

87. J. L. Carvalho-de-Souza, F. Bezanilla, Noncanonical mechanism of voltage sensor coupling to pore revealed by tandem dimers of Shaker. Nat. Commun. 10, 3584 (2019).

88. W. Wang, R. MacKinnon, Cryo-EM Structure of the Open Human Ether-a-go-go-Related K(+) Channel hERG. Cell 169, 422–430 e410 (2017).

89. X. Tao, R. MacKinnon, Cryo-EM structure of the KvAP channel reveals a non-domain-swapped voltage sensor topology. Elife 8, (2019).

90. C. H. Lee, R. MacKinnon, Structures of the Human HCN1 Hyperpolarization-Activated Channel. Cell 168, 111–120 e111 (2017).

91. J. R. Whicher, R. MacKinnon, Structure of the voltage-gated K(+) channel Eag1 reveals an alternative voltage sensing mechanism. Science 353, 664–669 (2016).

92. M. Li et al., Structure of a eukaryotic cyclic-nucleotide-gated channel. Nature 542, 60–65 (2017).

93. J. Shi, J. Cui, Intracellular Mg^2+^ Enhances the Function of Bk-Type Ca^2+^-Activated K^+^ Channels. The Journal of General Physiology 118, 589–606 (2001).

94. G. Gessner et al., Molecular mechanism of pharmacological activation of BK channels. Proceedings of the National Academy of Sciences 109, 3552–3557 (2012).

95. M. Yazdani et al., Aromatic interactions with membrane modulate human BK channel activation. Elife 9, e55571 (2020).

96. J. Lee et al., CHARMM-GUI input generator for NAMD, GROMACS, AMBER, OpenMM, and CHARMM/OpenMM simulations using the CHARMM36 additive force field. J. Chem. Theory Comput. 12, 405–413 (2016).

97. J. Huang et al., CHARMM36m: an improved force field for folded and intrinsically disordered proteins. Nat. Methods 14, 71–73 (2017).

98. J. B. Klauda et al., Update of the CHARMM all-atom additive force field for lipids: validation on six lipid types. J Phys Chem B 114, 7830–7843 (2010).

99. K. J. Bowers et al., paper presented at the Proceedings of the 2006 ACM/IEEE conference on Supercomputing, Tampa, Florida, 2006.

100. B. Hess, C. Kutzner, D. van der Spoel, E. Lindahl, GROMACS 4: algorithms for highly efficient, load-balanced, and scalable molecular simulation. J. Chem. Theory Comput. 4, 435–447 (2008).

101. M. J. Abraham et al., GROMACS: High performance molecular simulations through multi-level parallelism from laptops to supercomputers. SoftwareX 1–2, 19-25 (2015).

102. T. Darden, D. York, L. Pedersen, Particle mesh Ewald: An *N*-log (*N*) method for Ewald sums in large systems. The Journal of Chemical Physics 98, 10089 (1993).

103. J.-P. Ryckaert, G. Ciccotti, H. J. C. Berendsen, Numerical integration of the cartesian equations of motion of a system with constraints: molecular dynamics of n-alkanes. J. Comput. Phys. 23, 327–341 (1977).

104. S. Nosé, A unified formulation of the constant temperature molecular dynamics methods. The Journal of Chemical Physics 81, 511–519 (1984).

105. W. G. Hoover, Canonical dynamics: Equilibrium phase-space distributions. Phys. Rev. A 31, 1695–1697 (1985).

106. M. Parrinello, A. Rahman, Polymorphic transitions in single crystals: A new molecular dynamics method. J. Appl. Phys. 52, 7182–7190 (1981).

107. S. Jo, T. Kim, W. Im, Automated Builder and Database of Protein/Membrane Complexes for Molecular Dynamics Simulations. PLoS One 2, e880 (2007).

108. N. Michaud-Agrawal, E. J. Denning, T. B. Woolf, O. Beckstein, MDAnalysis: A toolkit for the analysis of molecular dynamics simulations. J. Comput. Chem. 32, 2319–2327 (2011).

109. O. S. Smart, J. G. Neduvelil, X. Wang, B. A. Wallace, M. S. P. Sansom, HOLE: A program for the analysis of the pore dimensions of ion channel structural models. J. Mol. Graphics Modell. 14, 354-& (1996).

110. A. Aksimentiev, K. Schulten, Imaging alpha-hemolysin with molecular dynamics: ionic conductance, osmotic permeability, and the electrostatic potential map. Biophys. J. 88, 3745–3761 (2005).

111. W. Humphrey, A. Dalke, K. Schulten, VMD: Visual molecular dynamics. Journal of Molecular Graphics 14, 33–38 (1996).

112. J. Eargle, Z. Luthey-Schulten, NetworkView: 3D display and analysis of protein.RNA interaction networks. Bioinformatics 28, 3000–3001 (2012).

113. R. W. Floyd, Algorithm 97: Shortest path. Commun. ACM 5, 345 (1962).

114. M. Girvan, M. E. Newman, Community structure in social and biological networks. Proc. Natl. Acad. Sci. U. S. A. 99, 7821–7826 (2002).

115. P. W. Kang et al., Calmodulin acts as a state-dependent switch to control a cardiac potassium channel opening. bioRxiv, 2020.2007.2004.187161 (2020).

116. A. M. Westerlund, O. Fleetwood, S. Perez-Conesa, L. Delemotte, Network analysis reveals how lipids and other cofactors influence membrane protein allostery. J. Chem. Phys. 153, (2020).

117. A. P. Dempster, N. M. Laird, D. B. Rubin, Maximum Likelihood from Incomplete Data Via the EM Algorithm. Journal of the Royal Statistical Society: Series B (Methodological) 39, 1–22 (1977).

