## Supplemental tables and figures for "Atomistic Simulation of Voltage Activation of a Truncated BK Channel"

#### Supplementary Tables

**Table S1. Summary of simulations.** The closed state structure is the fully equilibrated conformation derived from the $Ca^{2+}$-free Cryo-EM structures of the *ac*BK channel (PDB 5tji), and the activated state structure (*sim 5* and *6*) was taken from the last snapshot of *sim2b*. The P-loop/filter (T273 to D292) and C-terminus were harmonically restrained with a force constant of 0.5 $kcal.mol^{-1}.Å^{-2}$ in all production simulations. The 7 VSD charged residues examined in *sims 3-6* are D153, R167, D186, R207, R210, R213 and E219.

| Runs | Biased Potential | Initial State | Voltage (mV) | Platform | Simulation Lengths |
|---|---|---|---|---|---|
| *sim 1* | N/A | closed | 0 | Anton 2 | 2.0 μs |
| *sim 2* | N/A | closed | 750 | Anton 2 | 10 μs x 2 (a, b) |
| *sim 3* | N/A | resting | 0 | GPU Gromacs | 7 residues x 11 *λ* * 4 ns |
| *sim 4* | N/A | resting | 750 | GPU Gromacs | 7 residues x 11 *λ* x 4 ns |
| *sim 5* | N/A | activated | 0 | GPU Gromacs | 7 residues x 11 *λ* x 4 ns |
| *sim 6* | N/A | activated | 750 | GPU Gromacs | 7 residues x 11 *λ* x 4 ns |
| *sim 7* | N/A | Cryo-EM open state | 750 | GPU Gromacs | 0.4 μs x 1 |
| *sim 8* | SMD | closed | 0 | GPU Gromacs | 0.8 μs x 2* |
| *sim 9* | N/A | activated | 300 | GPU Gromacs | 1.0 μs x 3 |

* In *Sim 8*, the S5 and S6 became detached in two out of four replicas and the simulations were terminated before reaching the end of 0.8 μs.

#### Supplementary Movies

Due to file size restriction, movies can be found through the following link:
https://drive.google.com/drive/folders/1TMUQh6gJTL85V_L7vUOJbX8oB733mGXv

**Movie S1**: Movement of gating charges and TM helices S1-S6 during voltage activation of Core-MT BK channel (sim 2b). Key charges R210 (S4) and R213 (S4) are colored in blue, D153 (S2) and D182 (S3) in red. In addition, conserved S6 charges, E321 and E324, at the cytosolic entrance of the pore are shown in red sticks.

**Movie S2**: Movement of pore lining S6 helices (red cartoon) during voltage activation of Core-MT BK channel (sim 2b). The view is from the bottom (cytosolic side). The arrangement of S6 helices in the Ca2+-bound state of full-length BK channel is show in yellow cartoon for reference. E321 and E324 at the cytosolic entrance of the pore are shown in green sticks.

**Movie S3**: $K^+$ permeation events during the last 1 us of sim 2b, when the dilated pore is hydrated and conductive. The K+ ions are presented as spheres and colored by the atom index. Water molecules within the pore region are shown using red sticks.

### Supplementary Figures

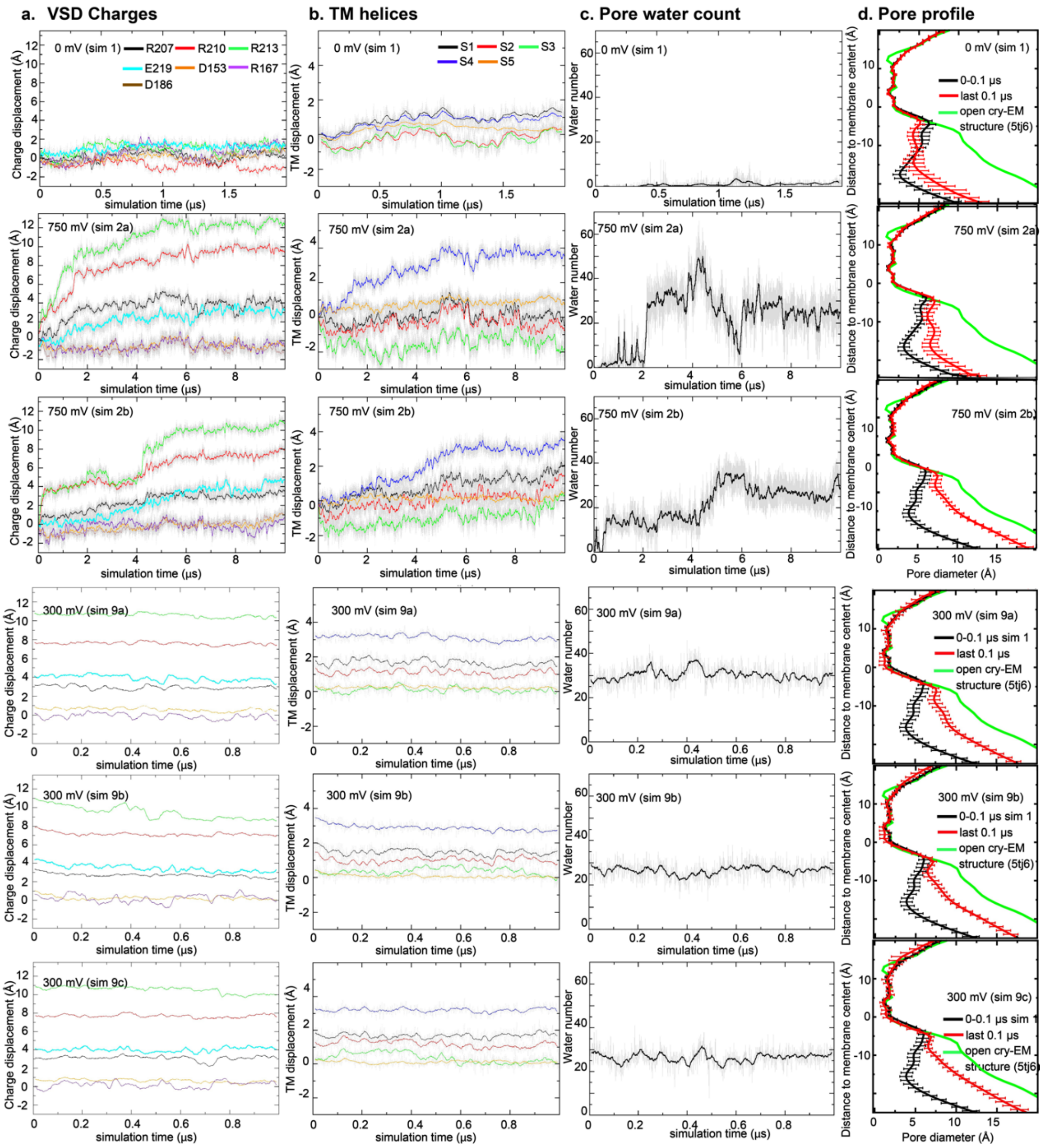


**Figure S1** The charged group z-displacement, z-displacement of the centers of mass of TM helices, number of pore waters, the averaged pore profiles during four Anton 2 simulations of Core-MT BK channels at 0 mV (row 1), 750 mV (rows 2-3) and 300 mV (rows 4-6) membrane voltages. See Methods for additional details of the simulation and analysis.

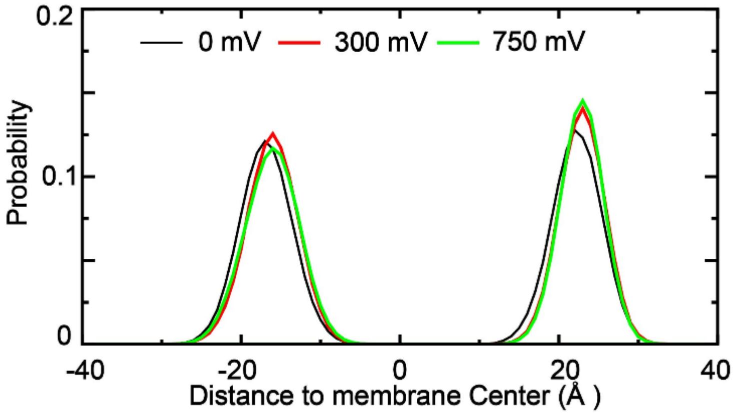


**Figure S2.** Distribution of lipid phosphor atoms under different membrane voltages. The distributions were derived from the last 500 ns of simulation in 0 mV (sim 1), 300 mV (sim 9) and 759 mV (sim 2a). Only the phosphate groups not within 12 Å of any protein atoms were selected for analysis. The average distances between phosphor atoms in the upper and lower leaflets are 45.6 Å, 47.0 Å and 47.0 Å at 0 mV, 300 mV and 750 mV, respectively. One-way ANOVA suggest that the difference in average membrane thickness under different voltages is not significant.

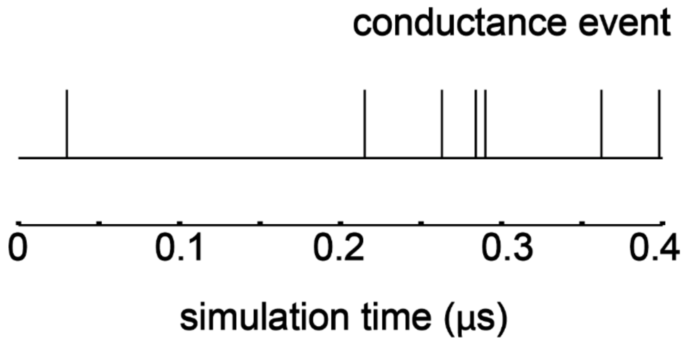


**Figure S3.** Conductance of the open state of Core-MT derived from the $Ca^{2+}$-bound full-length BK structure (PDB: 5tj6). The voltage was set at 750 mV and ion permeation events are shown as impulses. The conductance estimated from the second half of the trajectory is ~6 pS.

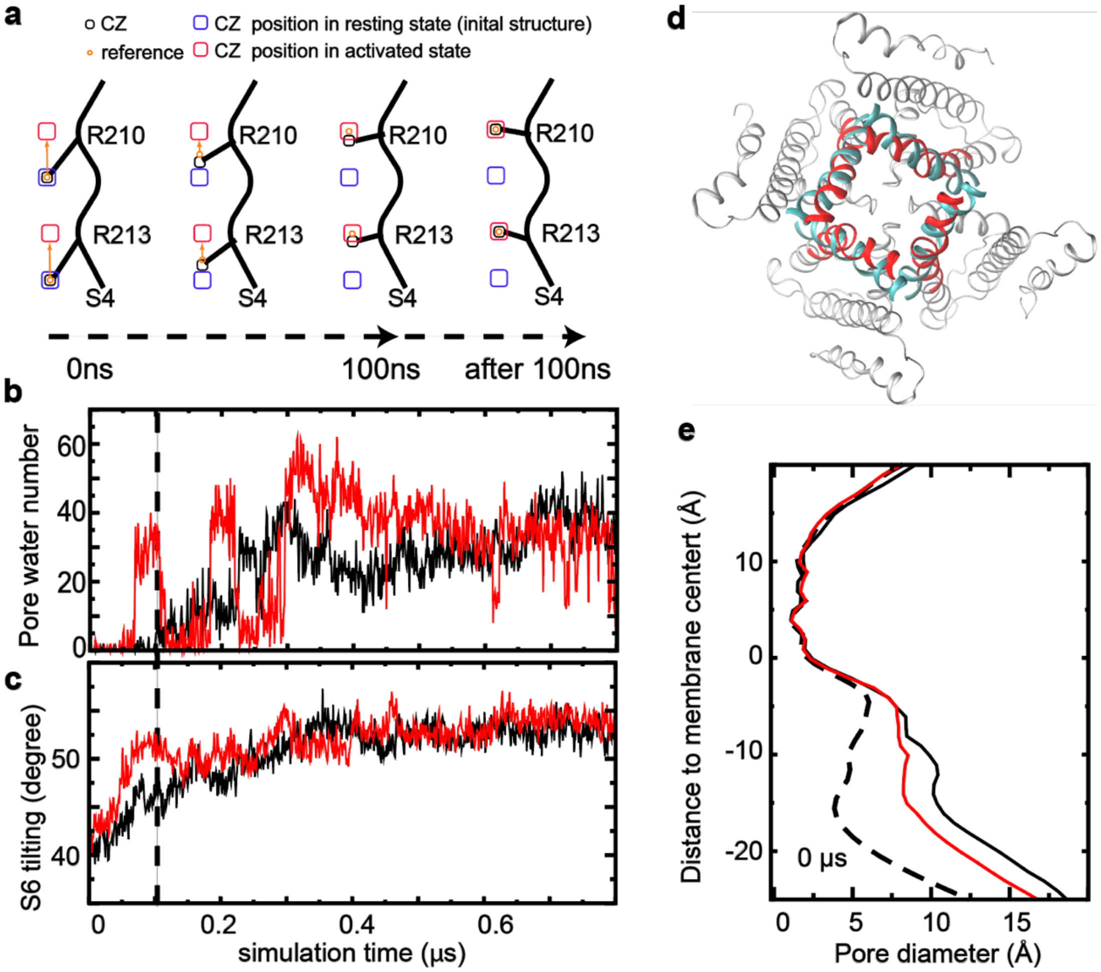


**Figure S4. Steered MD simulations of BK activation. a)** Illustration of the steered MD setup. Initially, the reference point (orange circle) is placed at the position of R210 or R213 guanidinium CZ atom in the resting state (black circle/blue rectangle). From 0-100 ns, the reference point moves along the z-axis (orange arrow) from the resting state position towards the activated state position (red rectangle). A harmonic positional restraint of 5 kcal/mol·Å², applied along the z-axis only, was used between the reference point and the CZ atom, steering the CZ atom to move along the z-axis with the reference. After 100 ns, the reference point remains fixed at the activated state position (red rectangle), allowing the VSD and the rest of the channel to respond to R210 and R213 movements. **b** and **c)** Numbers of pore water and S6 tilting angle as a function of simulation time during two of the steer MD simulations that lead to pore opening (black trace: replica 2; red trace: replica 4). **d)** Overlay of the pore structure at the end of replica 2 (red) and the open Cryo-EM structure (cyan). **e)** Pore profile at the end of replica 2 (red) in comparison to those from the initial state (dashed line) and the open Cryo-EM structure (black).

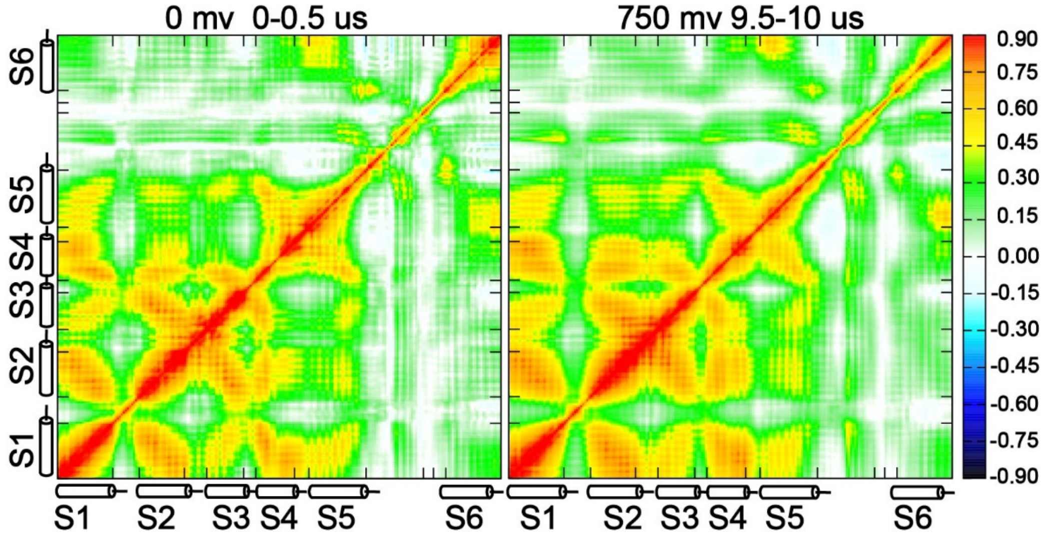


**Figure S5.** Covariance matrices of the Core-MT BK channel before (left) and after (right) voltage-induced activation (sim 2b).

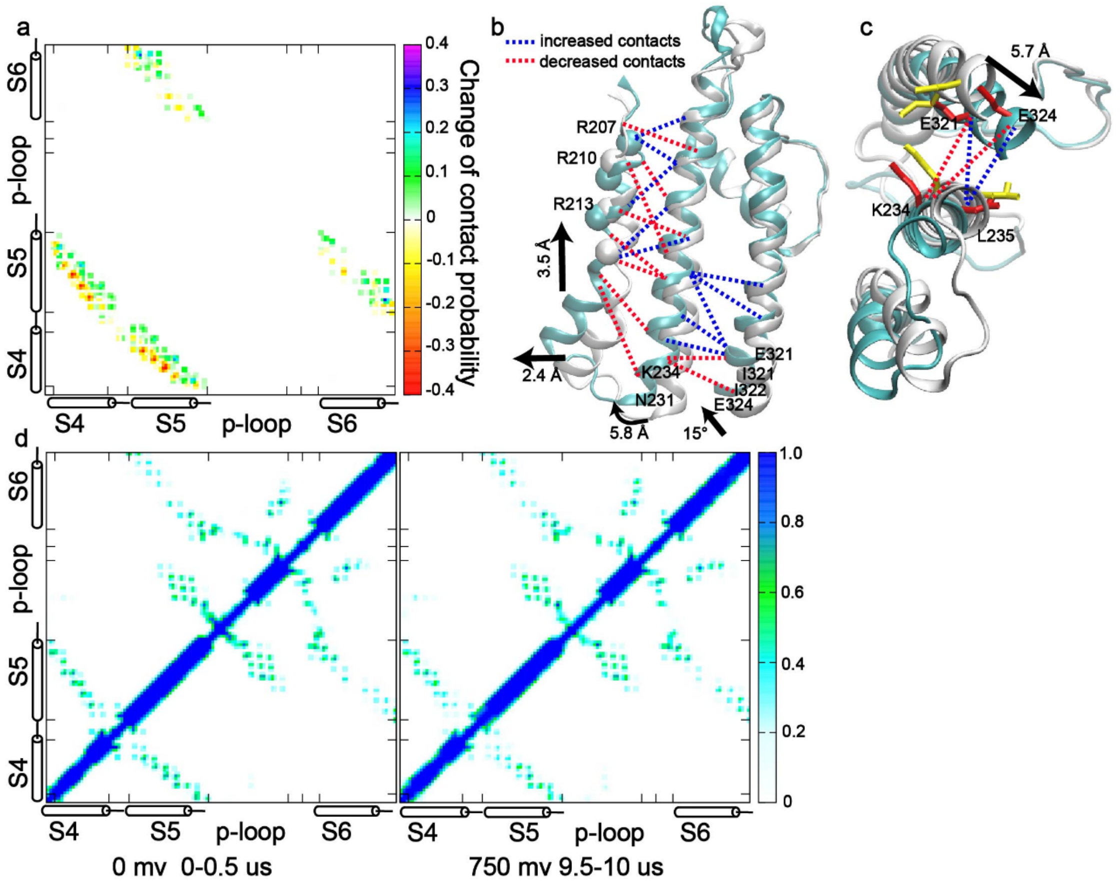


**Figure S6. Locking and concerted movements of S4, S5 and S6. a)**. The difference of residue-residue contact probability between the closed (sim 1, 0-0.5 μs) and activated state (sim 2b, 9.5-10 μs). **b)** Side and **c)** bottom view of the average activated (cyan) and closed (white) structures. The structures are aligned using the filter and P-loop. Dashed lines depict contacts where the probability increased (blue) or decreased (red) by more than 0.4 after activation. The Cα atoms of R207, R210, and R213 are represented as spheres. Residues (K234, L235, E321, and E324) involved in S6 bending are illustrated as yellow and red sticks for closed and activated states, respectively; **d)** Contact maps of S4-6 before (left) and after (right) voltage activation of Core-MT BK channels.

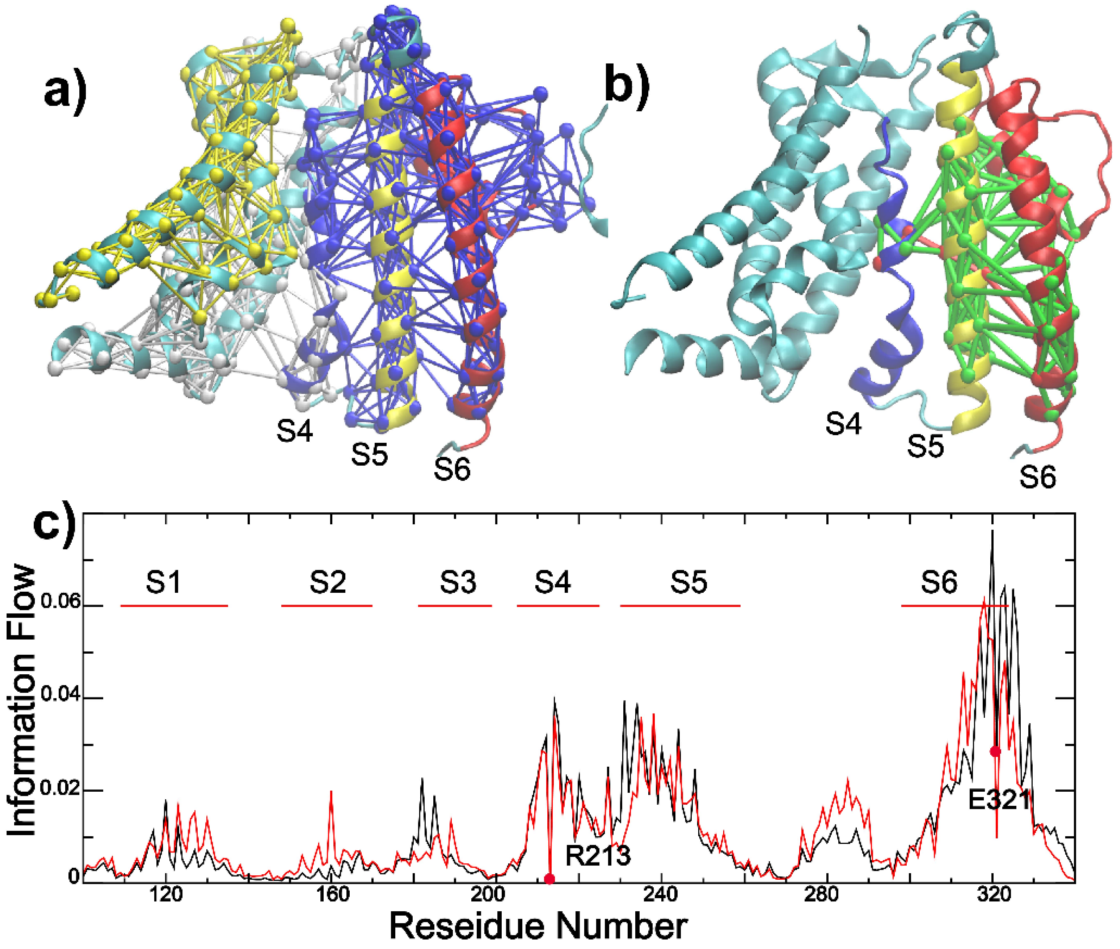


**Figure S7: Dynamic community, coupling pathways, and information flow of VSD-pore coupling derived from simulation of $Ca^{2+}$-bound structure.** All results were derived from the last 200 ns of *sim* 7 (Table S1). **a)** Dynamic community analysis showing that TM S4-6 are clustered into single tightly coupled community (blue network). The nodes (residues) and edges (contacts) are colored based on the community number. **b)** Optimal and suboptimal pathways of dynamic coupling between R213 (VSD S4) and E321 (pore-lining S6). All paths are colored green except for the optimal path, which is colored red. **c)** Information flow profile with R213 as the source and E321 as the sink node (labeled by red circle), respectively. Dynamic coupling analysis from last 500 ns of *sim 1* (closed state at 0 mV) is also show as reference (black trace).

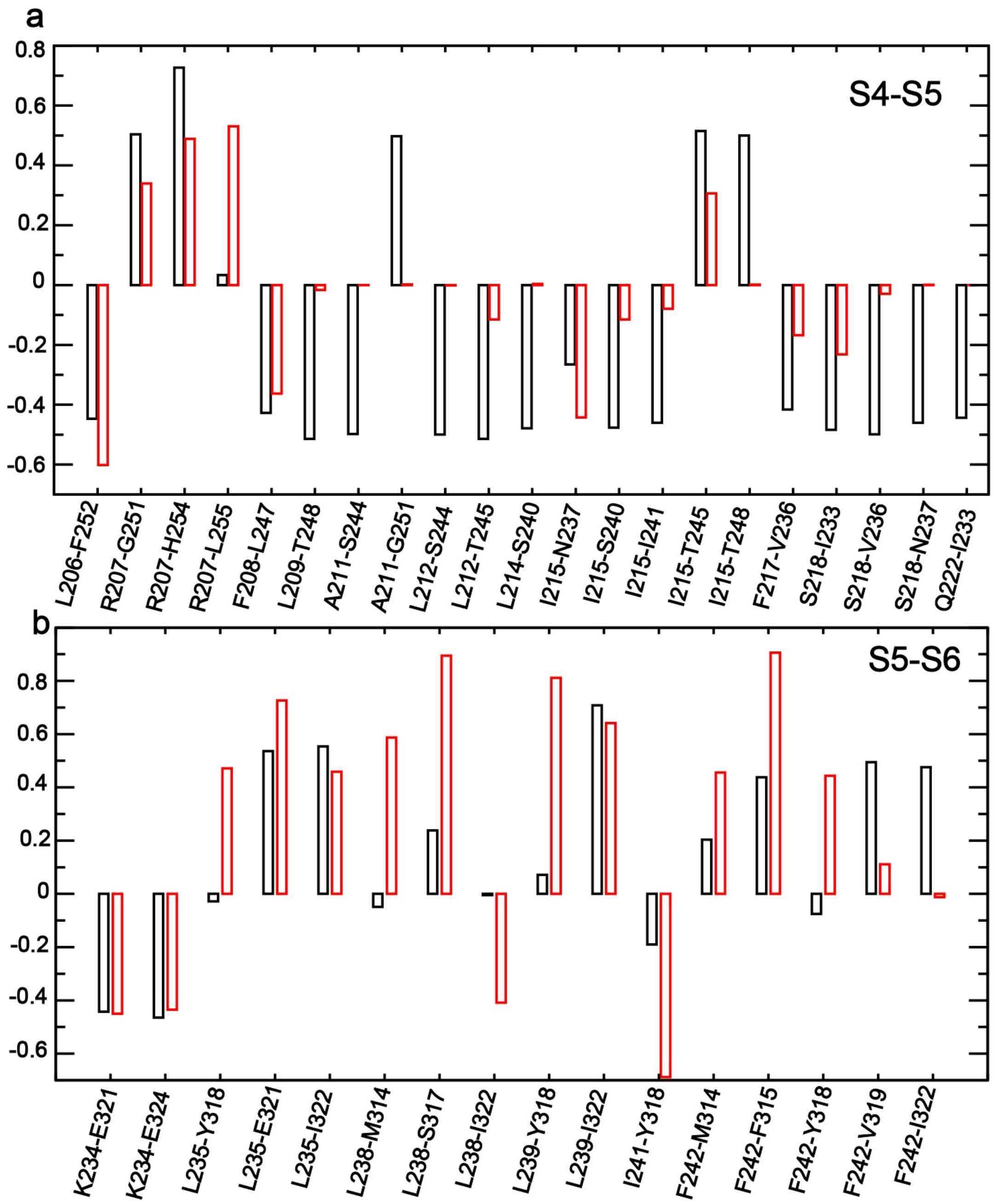


**Figure S8. Changes of residue-residue contact probability relative to the closed channel. a) S4-S5 interface, and b) S5-S6 interface.** The contact probabilities of the closed state were derived from *sim 1* (0–0.5 μs). Black bars show the changes observed in the voltage-driven activated state from *sim 2b* (9.5–10 μs), and red bars show the changes in the cryo-EM open state from *sim 7* (200–400 ns). Only residue pairs with a contact probability change of at least 0.4 in either *sim 2b* or *sim 7* are shown for clarity.
